# Gut Microbiota Fuel Aging Induced by Mitochondrial Dysfunction

**DOI:** 10.64898/2026.10.02.756159

**Authors:** Francesca Solagna, Daniel Martínez-Martínez, Norma Gabriela Zavala Torres, Annamaria Regina, Jonas Widder, Göksu Gökberk Kaya, Cosimo De Napoli, Schmidt Luisa, Ayesha Safoor, Jennifer van der Laan, Kaylee M. Morris, Susanne Brodesser, Aleksandra Trifunovic, Leonardo Nogara, Bert Blaauw, Marcus Krüger, Manolis Pasparakis, David E. Sanin, Mauro Corrado, Filipe Cabreiro

## Abstract

Aging is characterized by hallmarks whose mechanistic interdependencies remain poorly defined. Here, we investigate the bi-directional communication between gut microbiota and host mitochondria in the context of immune regulation, healthspan and lifespan. Using a mitochondrial dysfunction PolgA^mut^ mouse model of accelerated aging, we tested the role of mitochondrial dysfunction in modulating microbial homeostasis and aging. We find that impairment of mitochondrial function hastens the decline of microbiota structure over time and induces immune dysregulation and inflammation through metabolic reprogramming of the myeloid immune subset. Further, we find that gut microbiota drives a paradoxical energy state by reducing mitochondrial function systemically while simultaneously driving energy-intensive programs including immune remodelling and protein synthesis, through mTOR signaling. Thus, aging emerges from the progressive imbalance between microbiota-driven energetic demand and declining mitochondrial energy supply. We postulate that microbiota promote host early-life fitness, at the cost of amplifying metabolic stress and inflammation later in life.

**HIGHLIGHTS:**

- Host-microbiota homeostasis imposes an energetic cost that accelerates aging
- Dysbiosis and immune myeloid reprogramming are hallmarks of both natural and accelerated aging
- Microbiota diminish OXPHOS subunit expression and activity systemically with age
- Microbiota amplify mTOR signaling and metabolic stress in peripheral tissues

## INTRODUCTION

The intestine represents a uniquely demanding physiological interface: it sustains rapid epithelial renewal, harbors a large proportion of immune cells, and coexists with dense microbial communities. The latter are collectively referred to as the gut microbiota and represent a functional organ that has co-evolved with their hosts and is strongly shaped by both host factors, as well as environmental cues such as diet, medication and other lifestyle factors. This symbiotic relationship provides numerous advantages at different life stages, including but not limited to the fermentation of indigestible complex fibers, colonization resistance against pathogens, modulation of the immune system, and the production of essential vitamins ^1,2^. Furthermore, gut microbiome patterns and uniqueness within human populations reflect healthy aging and predict survival, hence gut microbiome changes occurring with age being used to develop aging clocks ^3–6^. Within this intricate framework, the causal contribution of the gut microbiome to shaping aging trajectories and driving organ-specific age-related pathologies remains mechanistically uncertain.

Aging is characterized by multiple interconnected hallmarks, including immune dysregulation, chronic low-grade inflammation, mitochondrial decline, and alterations in the gut microbiota ^7,8^. Over a century ago, Élie Metchnikoff postulated that microbial translocation across a compromised intestinal barrier promotes systemic inflammation and age-associated decline ^9^. This initial hypothesis anticipated the fundamental role of the gut and its microbes as important biological modulators of aging. Work in flies and mice partly validated this theory, showing that age-associated microbial dysbiosis promotes intestinal permeability, systemic inflammation and macrophage dysfunction ^10–12^. Fecal microbiota transplantation was previously shown to extend healthspan and lifespan in progeroid mice, supporting a causal contribution of the gut microbiota to aging-associated decline ^13^. Yet to-date, the interplay between dysfunctional hallmarks of aging remains vastly understudied ^10,12^. Mitochondria are key cellular organelles in this regard. Beyond energy generation through oxidative phosphorylation (OXPHOS), mitochondria act as signaling organelles that regulate redox balance, apoptosis, and immune pathways ^14^. The communication between microbes and mitochondria is a fascinating interaction, with several new findings revealing bacterial metabolites which can fine-tune mitochondrial dynamics, thereby influencing host metabolic fitness and longevity ^15–17^, but the reverse communication has not yet been studied. Importantly, mitochondrial function deteriorates progressively with age and is a defining feature of the aging process ^8^. In humans, mtDNA mutations accumulate markedly from approximately 60 years of age, driven primarily by replication-associated errors rather than oxidative stress. This replication-centric model of mutagenesis highlights intrinsic limitations in mtDNA maintenance and fidelity as key contributors to mitochondrial decline during aging ^18^. In mice, accumulation of mitochondrial DNA (mtDNA) mutations is causally linked to premature aging phenotypes in the PolgA^D257A^ mtDNA mutator (PolgA^mut^) mouse model that carries a missense mutation which drastically reduces 3′–5′ exonuclease activity required for proof-reading. PolgA^mut^ mice display progeroid phenotypes including alopecia, loss of body fat, sarcopenia, kyphosis, anemia, osteoporosis, cardiomyopathy, and a shortened lifespan ^19^.

Here, we explore the bi-directional interplay between gut microbiota and host mitochondria during aging. In particular, we examine the role of dysregulated mitochondrial function in a progeroid PolgA^mut^ mouse model and the consequences of mitochondrial interaction with the microbiota for systemic physiology and lifespan. First, we investigated whether PolgA^mut^ mitochondrial dysfunction-induced progeria caused a shift in gut microbial communities, and observed accelerated dysbiosis with a concomitant increase in systemic inflammation in an age- and genotype-dependent manner. Next, to address causality, we generate the first germ-free (GF) PolgA^mut^ mouse model, and demonstrated that absence of the microbiota markedly attenuated intestinal pathology and suppressed both local and systemic inflammation. Strikingly, microbial depletion mitigated the detrimental effects of mitochondrial dysfunction on organismal physiology, robustly extending healthspan and lifespan. Overall, our data show that, as mitochondrial efficiency declines, the cost of maintaining host-microbiota homeostasis shortens lifespan.

## RESULTS

### Mitochondrial dysfunction anticipates the age-dependent dysbiosis and inflammatory program observed in wild-type aged mice

To investigate the bi-directional interaction between the gut microbiota and host mitochondria, we first assessed the impact of mitochondrial deficiency on gut microbiota composition. We performed longitudinal shotgun metagenomics by comparing wild-type (WT) and PolgA^mut^ mutant mice at three different ages: 10 weeks, 26 weeks (coincident with pronounced weight loss the PolgA^mut^ genotype), and 38 weeks (prior to the onset of mortality) (Figure 1A). Beta-diversity analysis revealed a pronounced age-dependent change of the gut microbiota in PolgA^mut^ mice. Principal coordinate analysis (PCoA) using Jaccard distances (Figure 1B) showed a significant shift of gut microbial communities in the first dimension for the PolgA^mut^ mice over time, suggesting accelerated age-related dysbiosis. At the phylum level, we observed a significant expansion of Firmicutes (with a significant contribution of species belonging to the *Lactobacillus*, *Ligilactobacillus* and *Limosilactobacillus* genera (Figure S1A) and a decrease of Bacteroidetes (driven by a contribution of species from the *Muribaculum, Odoribacter* and *Cryptobacteroides* genera) with age in a genotype-specific manner (Figure 1C, Supplementary Table 1), consistent with animal microbiota dysbiotic signatures in WT aged animals ^4^. This age-related effect was further confirmed by applying Taylor’s law parameters ^20^ that show a loss of microbiota stability in a genotype x age-dependent manner (Figure 1D), with several species contributing to this relationship (Figure S1A) ^21^. Given the central role of short-chain fatty acids (SCFAs) in host-microbiota crosstalk and their potential role in aging ^22^, we interrogated the abundance of acetate, propionate, and butyrate in the host plasma (Figure S1B) and stool samples (Figure S1C). Collectively, these data indicate that the PolgA^mut^ model of accelerated aging drives remodelling of microbial community composition without significantly impacting overall SCFA production and its abundance in the host.

**Figure 1.**
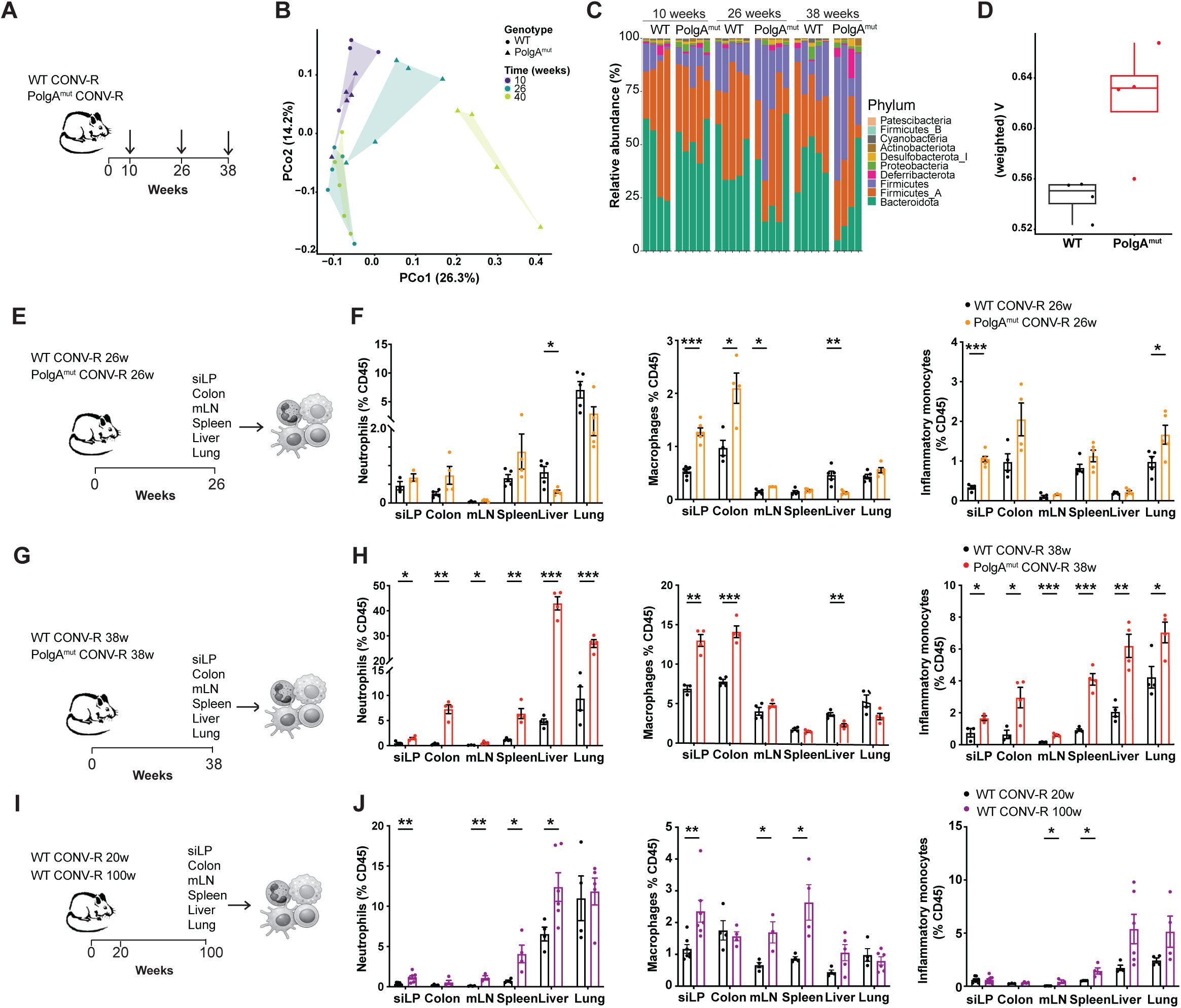
Mitochondrial dysfunction accelerates age-associated dysbiosis and inflammation. **(A)** Timeline of fecal sample collection from WT CONV-R and PolgA^mut^ CONV-R mice. **(B)** Principal coordinate analysis (PCoA) plot of the gut microbiota composition in stool samples collected from WT CONV-R and PolgA^mut^ CONV-R mice at the indicated time points. **(C)** Relative abundance of gut bacteria phyla in stool samples collected from WT CONV-R and PolgA^mut^ CONV-R mice at the indicated time points. **(D)** Assessment of microbiota stability by Taylor’s law analysis. **(E)** Schematic of multi-organ immunophenotyping of 26-week-old WT CONV-R and PolgA^mut^ CONV-R mice. **(F)** Frequencies of tissue-infiltrating neutrophils, classic macrophages and inflammatory monocytes in the indicated organs of 26-week-old WT CONV-R and PolgA^mut^ CONV-R mice. **(G)** Schematic of multi-organ immunophenotyping of 38-week-old WT CONV-R and PolgA^mut^ CONV-R mice. **(H)** Frequencies of tissue-infiltrating neutrophils, classic macrophages and inflammatory monocytes in the indicated organs of 38-week-old WT CONV-R and PolgA^mut^ CONV-R mice. **(I)** Schematic of multi-organ immunophenotyping of 20- and 100-week-old WT CONV-R mice. **(J)** Frequencies of tissue-infiltrating neutrophils, classic macrophages and inflammatory monocytes in the indicated organs of 20- and 100-week-old WT CONV-R mice. Data shown as mean ±SEM of three independent experiments unless indicated. Statistical comparisons between two groups were calculated by using an unpaired two-tailed Student’s t-test. *p < 0.05; **p < 0.01; ***p < 0.001.

Given the causal relationship of microbiota with immune regulation, we isolated immune cells from the small intestine lamina propria (siLP), colon, mesenteric lymph nodes (mLN), spleen, liver and lung of WT and PolgA^mut^ mice at 26 and 38 weeks of age (Figure 1E,G). We found an accumulation of myeloid cells in PolgA^mut^ mice (neutrophils, dendritic cells, macrophages and inflammatory monocytes), which appeared primarily in the colon and siLP of 26-week-old animals (Figure 1F, S1D). This phenotype worsened progressively over time, with increasing myeloid cell infiltration in the intestine and colon, also extending to peripheral tissues such as the spleen, liver and lung at 38 weeks of age (Figure 1H, S1D). Together with evidence from other immune dysfunction and aging models ^23^, these findings suggest that the intestine may provide a permissive environment for the onset of inflammation during aging, which subsequently extends systemically. Notably, the lymphoid cell compartment did not show major differences between WT and PolgA^mut^ mice, or change significantly over time, particularly the B cell and conventional TCRβ^+^ T cell subsets. However, at 38 weeks of age, PolgA^mut^ mice exhibited lower frequencies of CD4⁺ and CD8⁺ T cells in peripheral tissues, including the spleen, liver, and lung (Figure S1D). Similar findings were obtained in cohorts of young (20 weeks) and aged (100 weeks) WT mice (Figure 1I). WT aged mice exhibited increased myeloid cell-driven inflammation, evidenced by the recruitment and accumulation of neutrophils, dendritic cells, macrophages, and inflammatory monocytes (Figure 1J), as well as lower frequencies of CD4⁺ and CD8⁺ T cells (Figure S1E). Overall, these data suggest that mitochondrial dysfunction drives accelerated microbial dysbiosis and associated immune-related inflammation mimicking phenotypes observed with chronological aging in WT animals.

### Absence of gut microbiota extends lifespan and healthspan in a mitochondrial deficient model

Next, to test whether commensal gut microbes are causally linked to inflammaging in an accelerated model of aging, we generated germ-free (GF) PolgA^mut^ mice. Confirmation of GF status was performed at 38 weeks of age by measuring SCFAs in stool (Figure S1C). In the absence of microbes, PolgA^mut^ mice exhibited a markedly improved phenotype compared with conventionally raised (CONV-R) counterparts, including preserved body weight with age and a striking extension of lifespan (Figure 2A-C). Building on these findings, we next assessed the contribution of the microbiota to systemic inflammation by quantifying 32 serum cytokines, chemokines and growth factors in 38-week-old WT and PolgA^mut^ mice under CONV-R and GF conditions. PolgA^mut^ CONV-R mice displayed a pronounced hyper-inflammatory phenotype, characterized by significantly elevated levels of pro-inflammatory cytokines, including IL-6, IFN-γ, IL-12 and IL-7, as well as the chemokine MIP-1α, compared with both WT CONV-R and PolgA^mut^ GF cohorts (Figure 2D, S2A). This pro-inflammatory signature was largely absent in PolgA^mut^ GF mice, indicating that microbiota are a necessary trigger for systemic inflammation. Interestingly, levels of the potent anti-inflammatory cytokine IL-10 were increased exclusively in PolgA^mut^ CONV-R mice, likely indicating compensatory hyper-secretion. Elevated levels of lipopolysaccharide-binding protein (LBP) in PolgA^mut^ CONV-R mice further suggested that the inflammatory state was tied with compromised intestinal barrier integrity (Figure 2E). To determine whether intestinal tissue integrity was maintained given this cytokine profile, we performed histological analysis of the small intestine and colon. At the histological level, the colon of 38-week-old PolgA^mut^ CONV-R mice showed focal epithelial hyperplasia, increased immune cell infiltration in the lamina propria, mild epithelial erosion, and decreased goblet cell numbers in the transverse and proximal regions, compared with WT CONV-R and PolgA^mut^ GF cohorts. TUNEL staining revealed increased numbers of dying cells in the colons of PolgA^mut^ CONV-R mice. These alterations were largely absent in PolgA^mut^ mice maintained under GF conditions (Figure 2F, S2B). The small intestine of 38-week-old PolgA^mut^ CONV-R mice similarly displayed perturbed architecture, with blunted villi, decreased goblet and Paneth cell numbers, and increased cell death relative to WT CONV-R and PolgA^mut^ GF mice (Figure 2G). These alterations were also significantly attenuated in the absence of microbiota, as shown by quantitative assessment of pathology scores (Figure 2H). Overall, our accelerated aging model reveals that mitochondrial dysfunction establishes a baseline state of vulnerability, with the gut microbiota acting as a catalyst of systemic inflammation and aging-related phenotypes.

**Figure 2.**
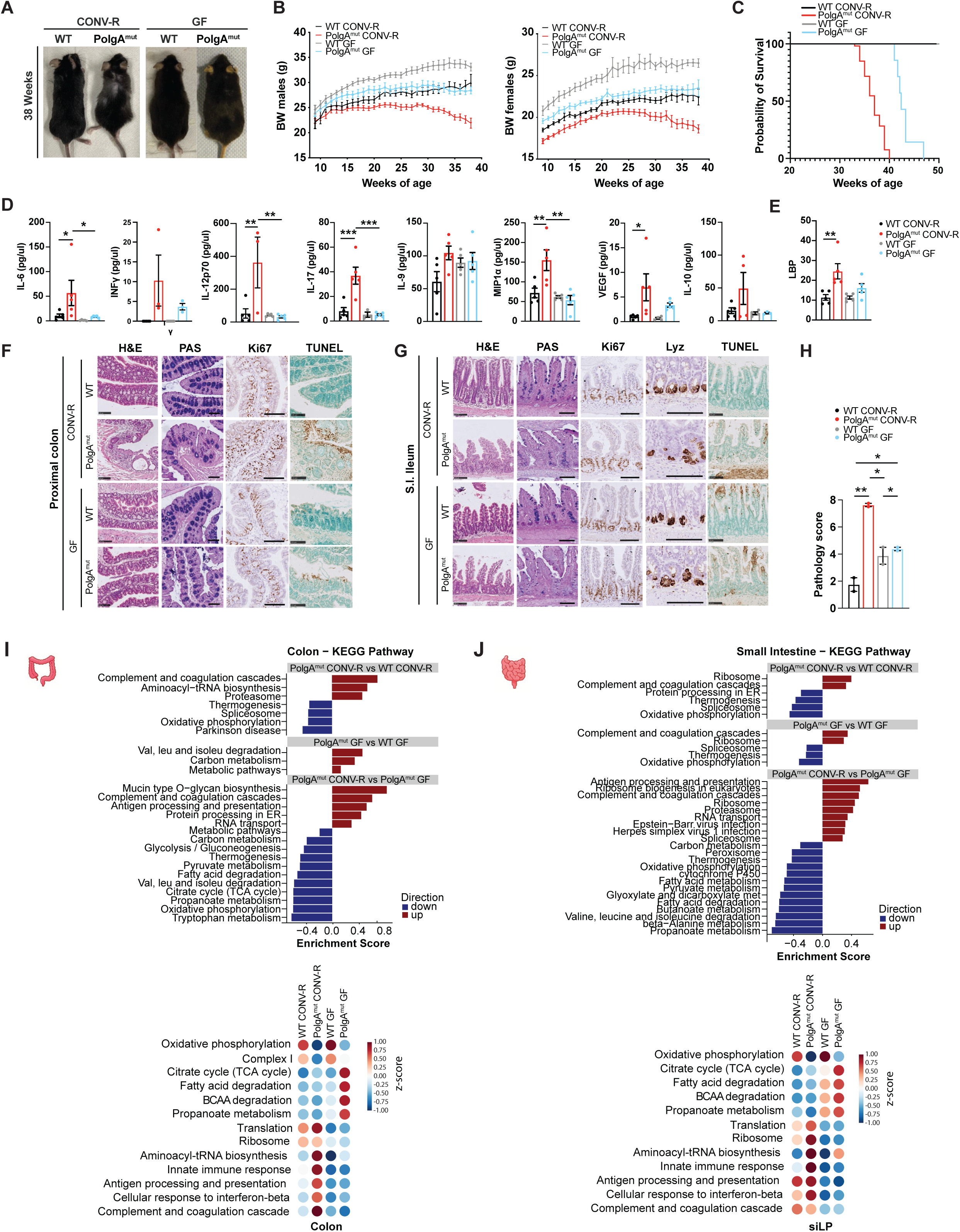
Absence of gut microbiota extends lifespan and healthspan in a mitochondrial deficient mouse model. **(A)** Representative pictures of 38-week-old CONV-R and GF WT and PolgA^mut^ mice. **(B)** Body weight curve of CONV-R WT (n=12), CONV-R PolgA^mut^ (n=9), GF WT (n=9), GF PolgA^mut^ (n=12) male mice. Body weight curve of CONV-R WT (n=12), CONV-R PolgA^mut^ (n=13), GF WT (n=10), GF PolgA^mut^ (n=6) female mice. **(C)** Kaplan-Meier survival curves of CONV-R WT (n=12), CONV-R PolgA^mut^ (n=53), GF WT (n=12), GF PolgA^mut^ (n=7) male and female mice. **(D)** Serum concentration levels of the indicated cytokines in 38-week-old CONV-R and GF WT and PolgA^mut^ mice. **(E)** Serum concentration levels of lipopolysaccharide-binding protein (LBP) in 38-week-old CONV-R and GF WT and PolgA^mut^ mice. **(F)** Representative images of proximal colon sections from 38-week-old CONV-R and GF WT and PolgA^mut^ mice stained with H&E, PAS, Ki67 and TUNEL staining. **(G)** Representative images of small intestine (ileum) sections from 38-week-old CONV-R and GF WT and PolgA^mut^ mice stained with H&E, PAS, Ki67, Lyz and TUNEL staining. **(H)** Pathology score of small intestine and colon of 38-weeks-old CONV-R and GF WT and PolgA^mut^ mice. **(I)** Pathway enrichment analysis of proteomic profile of colon samples from 38-week-old CONV-R and GF WT and PolgA^mut^ mice. **(J)** Pathway enrichment analysis of proteomic profile of small intestine samples from 38-week-old CONV-R and GF WT and PolgA^mut^ mice. Data shown as mean ±SEM of three independent experiments unless indicated. Statistical significance was determined using two-way ANOVA with Tukey’s correction for multiple comparisons; *p < 0.05, **p < 0.01, ***p < 0.001.

### Microbiota intensifies metabolic and immune proteome remodelling in PolgA ^mut^ mice

To define the impact of gut microbiota on mitochondrial deficiency in the colon and small intestine (ileum), we performed quantitative proteomics in WT and PolgA^mut^ mice under CONV-R and GF conditions at 38 weeks of age. Unbiased pathway enrichment analysis revealed consistent metabolic and immune alterations in both tissues across the different experimental conditions (Figure 2I-J, Supplemental Table 2). As expected, PolgA^mut^ CONV-R mice exhibited marked depletion of proteins involved in OXPHOS, including multiple Complex I (NDUF) and Complex IV (COX) subunits, consistent with decreased respiratory capacity, a hallmark of WT aging ^24^. Strikingly, expression of mitochondrial subunits was increased in both WT and PolgA^mut^ animals in the absence of gut microbes, suggesting a generalized non-genotype specific role played by the microbiota. In particular, defects observed in the PolgA^mut^ CONV-R animals were significantly attenuated under GF conditions, indicating that the presence of microbiota exacerbated the disruption of the mitochondrial proteome in tissues of the gastrointestinal tract (Figure 2I,J, S2D,E). Consistent with the impact on mitochondrial energy metabolism, many related metabolic pathways, such as TCA cycle, fatty acid degradation, branched-chain amino acid (BCAA) degradation and propanoate metabolism, were strongly upregulated in PolgA^mut^ GF mice. Pairwise comparisons of KEGG-enriched functions in a genotype x husbandry-dependent manner revealed these functions as equally important for explaining the phenotypes of proximal tissues such as the colon and small intestine, as well as distal tissues such as the liver. Furthermore, it highlighted the systemic impact of microbiota-mitochondria communication and the key role played by gut microbiota in regulating OXPHOS and functionally related pathways in PolgA^mut^ mice (Figure S2C). In contrast, pathways related to protein synthesis and processing (i.e. ribosome, translation, aminoacyl-tRNA biosynthesis, and proteasome) and immune-associated pathways (i.e. complement and coagulation cascades, antigen processing and presentation, and innate immune signaling), were among the most significantly enriched signatures in PolgA^mut^ CONV-R mice (Figure 2I,J S2D,E). Activation of these pathways was markedly blunted under GF conditions, especially in the colon. Altogether, these findings reveal a paradoxical state in which mitochondrial genome instability decreases ATP-generating capacity while simultaneously driving ATP-consuming protein synthesis and immune programs. The microbiota amplifies this vicious cycle, increasing the metabolic reprogramming associated with mitochondrial dysfunction and potentially accelerating functional decline with age.

### Absence of microbiota improves healthspan and muscle function through ameliorated mitochondrial function and decreased mTOR signaling

To determine how the absence of microbiota affects systemic energy metabolism in PolgA^mut^ mice, we examined skeletal muscle, a major regulator of whole-body metabolic homeostasis due to increased metabolic demand. Morphological analysis showed no clear signs of myofiber degeneration such as centrally nucleated fibers or associated-inflammation, but a decrease in fiber size was observed in PolgA^mut^ CONV-R compared to WT (Figure 3A). In contrast, GF PolgA^mut^ mice preserved fiber morphology (Figure 3A). Quantification of fiber cross-sectional area (CSA) confirmed a significant leftward shift of fiber size distribution in PolgA^mut^ CONV-R muscle, indicating a higher number of smaller fibers, whereas the GF condition rescued this defect to WT GF levels (Figure 3B). To assess muscle fiber function, we measured force production in isolated skinned fibers. Both CSA (Figure 3C) and tension (Figure 3D) were decreased in PolgA^mut^ CONV-R mice and were partially restored in PolgA^mut^ GF mice. Next, we performed quantitative proteomics in skeletal muscle from WT and PolgA^mut^ mice under CONV-R and GF husbandry to capture the molecular profiles underlying these changes. Hierarchical clustering of z-score–normalized protein with a significant interaction (P value<0.001) (n=486 proteins) identified six distinct clusters (C1-6) (Figure S3A, Supplemental Table 3). Notably, KEGG pathway enrichment analysis showed enrichment for pathways linked to energy-related metabolism, and to muscle contraction and fiber-type regulation. We observed a shift towards a slow-fiber program consistent with aged muscle and altered abundance of actin cytoskeleton-associated proteins in PolgA^mut^ CONV-R mice, which were rescued under GF conditions (Figure 3E,F, S3B). Furthermore, and consistent with the systemic effect of microbiota reported above, PolgA^mut^ CONV-R mice showed decreased expression of several proteins associated with mitochondrial electron transport chain complexes compared with WT CONV-R mice. In contrast, proteins involved in fatty acid metabolism and fatty acid oxidation (FAO) were increased in PolgA^mut^ CONV-R mice. This suggests that PolgA^mut^ mice shift towards greater lipid utilization, potentially as a response to impaired mitochondrial function. Changes in glycolytic and BCAA degradation proteins further support the presence of widespread metabolic adaptation in PolgA^mut^ CONV-R muscle that is partially reverted in PolgA^mut^ GF mice.

**Figure 3.**
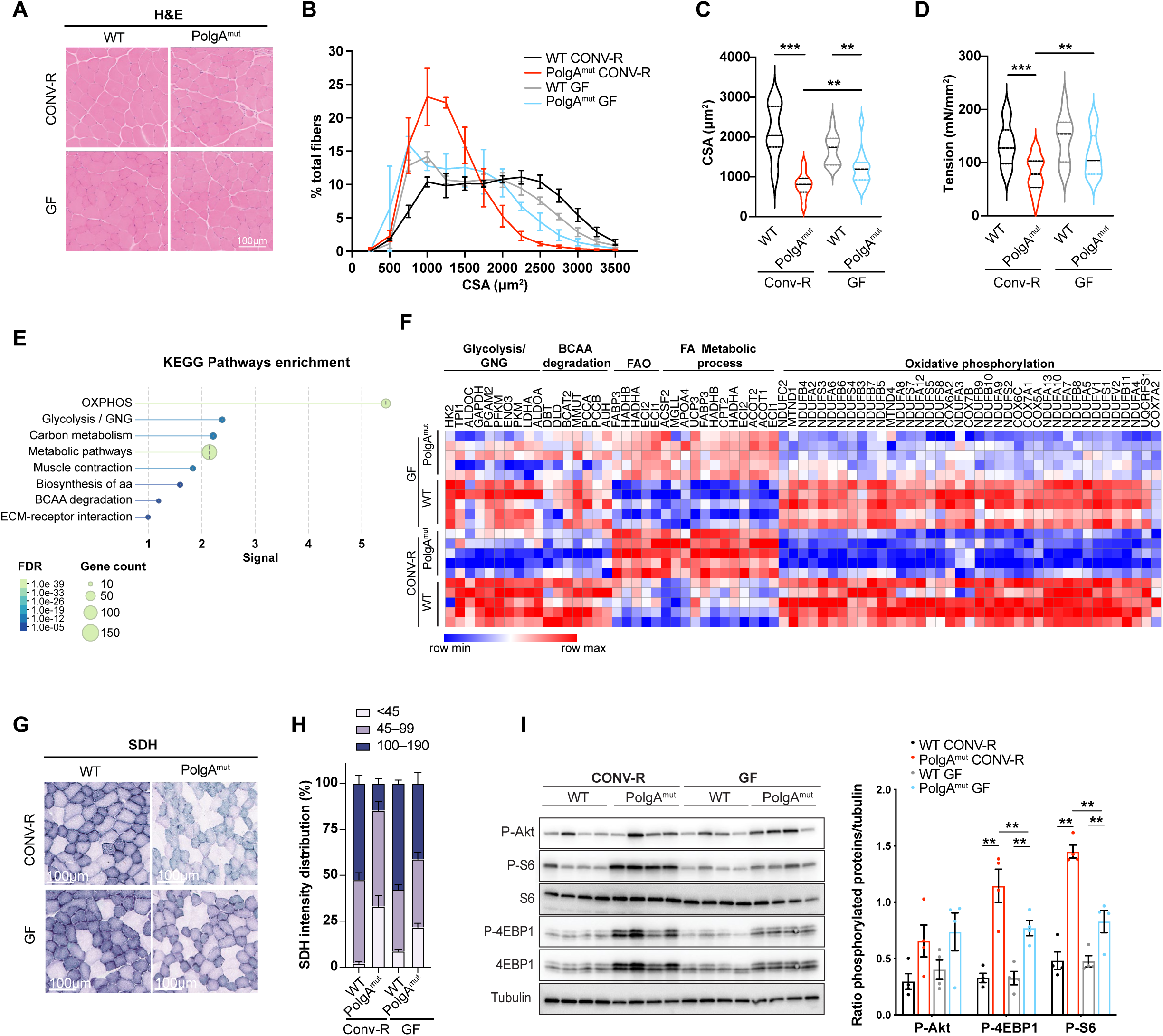
Absence of microbiota improves healthspan and muscle function through mTOR signaling. **(A)** Representative haematoxylin and eosin (H&E) staining of muscle from CONV-R and GF WT and PolgA^mut^ mice. Scale bars, 100 μm. **(B)** Frequency distribution of cross-sectional areas (CSA, μm^2^) in gastrocnemius (GC) of CONV-R and GF WT and PolgA^mut^ mice (n=3 mice per group) **(C)** Average cross-sectional area of skinned fibers from EDL muscle of CONV-R and GF WT and PolgA^mut^ mice (n=3 mice per group). **(D)** Maximal isometric tension produced by skinned fibers from EDL muscle of CONV-R and GF WT and PolgA^mut^ mice (n=3 mice per group, total fibers 12-16). **(E)** KEGG pathway enrichment analysis of hierarchical clustering of z-score–normalized protein with a significant interaction (p value<0.001) (n=486 proteins) of muscle from CONV-R and GF WT and PolgA^mut^ mice. **(F)** Heatmap of metabolic proteins involved in glycolysis/gluconeogenesis (GNG), BCAA degradation, FAO, fatty acid metabolic processes, and OXPHOS across experimental groups. **(G)** Representative succinate dehydrogenase (SDH) staining of muscle from CONV-R and GF WT and PolgA^mut^ mice. Scale bars, 100 μm. **(H)** Quantification of SDH-positive fiber populations classified by staining intensity. **(I)** Total protein extracts from muscle of 38-week-old CONV-R and GF WT and PolgA^mut^ mice, immunoblotted with the indicated antibodies. Data shown as mean ±SEM. Statistical significance was determined using two-way ANOVA with Tukey’s correction for multiple comparisons; *p < 0.05, **p < 0.01, ***p < 0.001.

Histological measurements of succinate dehydrogenase (SDH) activity in muscle tissue further confirmed the significant rescue of oxidative capacity by GF conditions in both WT and mitochondrial deficient PolgA^mut^ mice (Figure 3G, H). Scatter plots of SDH intensity versus fiber CSA showed a consistent inverse relationship across all groups. In particular, we observed a higher proportion of small fibers exhibiting lower SDH staining intensity in PolgA^mut^ CONV-R mice. This alteration was rescued in PolgA^mut^ GF mice, where the distribution of SDH intensity relative to CSA more closely resembled that of WT animals (Figure S3C). Emerging evidence indicates that age-dependent hyperactivation of mTOR in skeletal muscle may be implicated in sarcopenia and functional decline ^25,26^. Our data demonstrate that WT CONV-R mice at 100 weeks of age exhibit decreased CSA, reflected by a significant leftward shift in fiber size distribution relative to 20-week-old mice (Figure S3D), together with robust activation of the mTOR pathway, evidenced by increased phosphorylation of AKT (Ser473) and ribosomal protein S6 (rpS6) (Ser240/244), and inhibition of eIF4E-binding protein 1 (4EBP1) (Thr37/46) (Figure S3E). Interestingly, the hyperactivation of the mTOR pathway is attenuated in aged GF mice compared with CONV-R mice. Similarly to aged mice, we observed a significantly increased phosphorylation rpS6 (Ser240/244) and 4EBP1(Thr37/46) but not AKT (Ser473) in skeletal muscle of PolgA^mut^ CONV-R compared to control, consistent with activation of mTORC1 activity. Importantly, this hyperactivation was significantly decreased in the GF condition (Figure 3I). Altogether, these data indicate that the gut microbiota shape the metabolism of energy-demanding peripheral tissues in mitochondrial deficient mice and activates mTOR.

### Gut microbiota promotes inflammatory reprogramming of γδ T cells and myeloid cells in PolgA^mut^ mice

To test whether the ameliorated gut and systemic phenotypes in PolgA^mut^ GF mice could be traced back to an improved immune profile in the gut, we analysed the composition of the immune infiltrate in the siLP, colon, mLN and spleen of WT and PolgA^mut^ mice under CONV-R and GF conditions. GF husbandry strongly decreased the accumulation of myeloid cells observed in PolgA^mut^ CONV-R mice, especially neutrophils and inflammatory monocytes (Figure 4A, S4A). To define the composition and transcriptional profile of the immune infiltrate in an unbiased fashion, CD45^+^ cells sorted from small intestine and colonic lamina propria were analyzed by single-cell RNA sequencing (scRNAseq). Cell-type annotation using scGate ^27^ identified 24 transcriptionally distinct clusters covering the main immune cell subsets (Figure 4B, Supplemental Table 4). Comparative analysis revealed genotype-dependent remodeling of the myeloid compartment, with increased frequencies of cDC1 (cluster 6), cDC2 (cluster 7), macrophages (cluster 8), and pDCs (cluster 11) in PolgA^mut^ mice (Figure 4B,C, S4B). We also observed an increase of unconventional γδ T cells (cluster 15) in PolgA^mut^ CONV-R mice (Figure 4B,C, S4B). These cells are usually enriched at mucosal sites and act as rapid responders to microbial signals and tissue stress. In PolgA^mut^ CONV-R mice, γδ T cells showed a highly dysfunctional phenotype characterized by high expression of genes encoding for pro-inflammatory cytokines and exhaustion markers (*Pdcd1*, *Havcr2*, *Ctla4*, *Ly6a*, *Ccr5*, *Klrc1* and *Il17a*) (Figure 4D). Similarly, macrophages (cluster 8) presented higher expression of inflammatory genes, including *Il1b*, *Il1rn*, *Nfkbia*, *Ccrl2*, *Cd74*, *H2-Eb1*, *Fcer1g*, and *Tnfaip3*, more significantly in the colon of PolgA^mut^ CONV-R mice (Figure 4E). Notably, microbial status selectively affected both γδ T cells (cluster 15) and macrophages (cluster 8), which were significantly decreased in PolgA^mut^ GF mice compared to CONV-R conditions (Figure 4C, S4B). These findings suggest a compartment-specific role for these cells, with γδ T cells acting primarily in the siLP and macrophages in the colon. Pathway enrichment analysis showed upregulation of exhaustion and inflammatory programs in PolgA^mut^ CONV-R γδ T cells and macrophages. Pathways linked to DNA sensing, cellular senescence, apoptosis, and OXPHOS were also enriched in CONV-R conditions, suggesting activation of stress-response and metabolic rewiring. These signatures were markedly attenuated under GF conditions (Figure 4D,E). Metabolic pathways, including glycolysis/gluconeogenesis and TCA/citrate cycle, also displayed genotype- and microbiota-dependent modulation, suggesting a coordinated immunometabolic reprogramming in PolgA^mut^ γδ T cells and macrophages in the colon (Figure 4D,E). Given the elevated DNA sensing score across myeloid populations and the established activation of the mitochondrial (mt)DNA release-dependent Type I Interferon signature in PolgA^mut^ mice ^28^, we next examined the contribution of the cGAS-STING axis to the inflammatory phenotype by crossing PolgA^mut^ mice with STING-deficient animals. Genetic ablation of STING failed to rescue the intestinal immune phenotype with persisting neutrophil and inflammatory monocyte accumulation (Figure S4C). Altogether, these findings indicate that the inflammatory phenotype observed in PolgA^mut^ mice is largely STING-independent and instead, gut microbiota-dependent.

**Figure 4.**
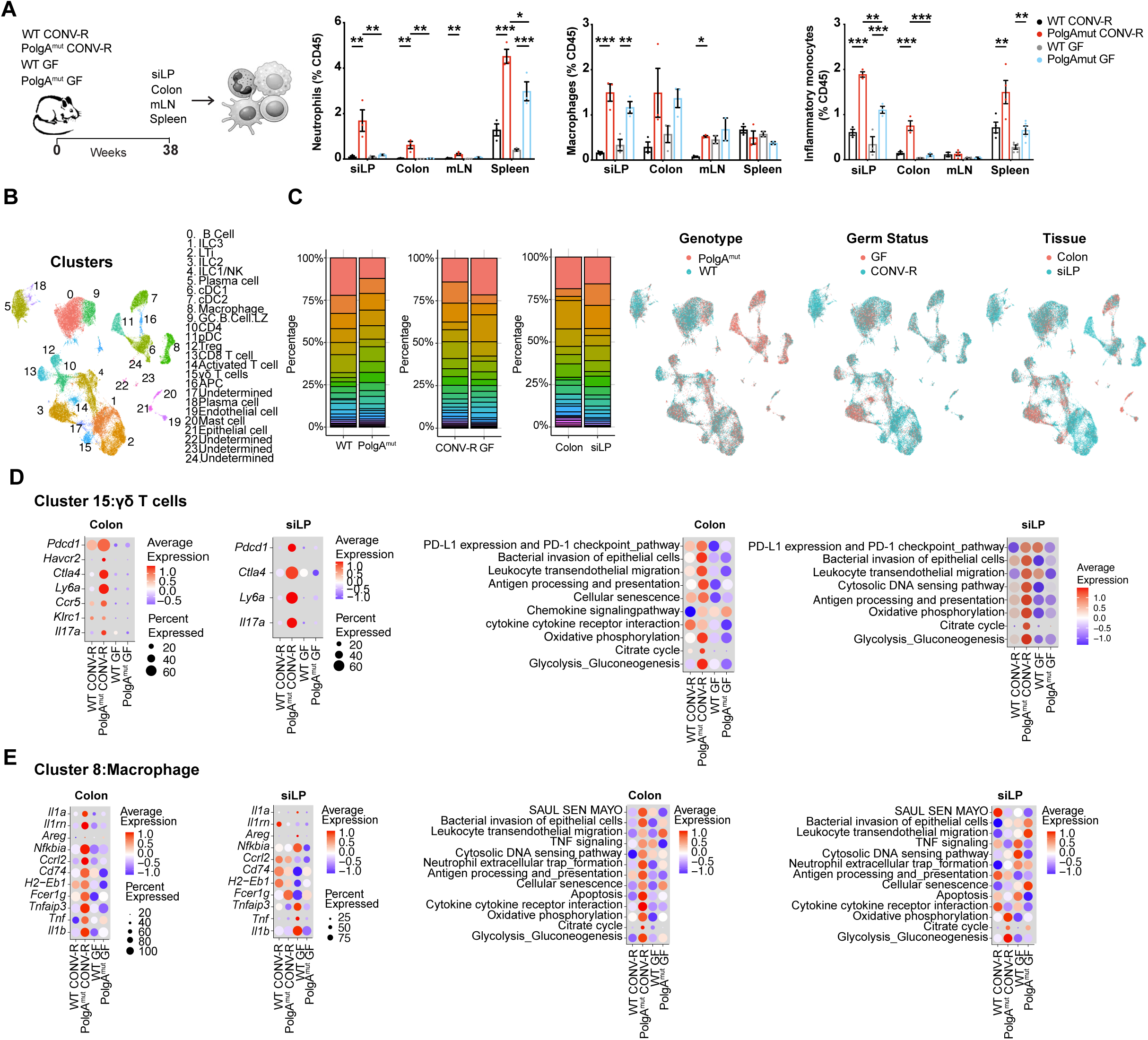
Gut microbiota promote inflammatory reprogramming of γδ T cells and myeloid cells in PolgA^mut^ mice. **(A)** Multi-organ immunophenotyping of 38-week-old CONV-R and GF WT and PolgA^mut^ mice with frequencies of tissue-infiltrating neutrophils, classic macrophages and inflammatory monocytes. **(B)** UMAP visualization of the scRNA-seq data showing the distribution of clusters and cell types identified in CD45^+^ cells sorted from the colon and small intestine of CONV-R and GF WT and PolgA^mut^ mice. **(C)** Frequencies of annotated cell clusters in CD45^+^ cells sorted from the colon and small intestine of CONV-R and GF WT and PolgA^mut^ mice (left). UMAP visualization distribution per genotype, germ status and tissue (right). **(D)** Gene expression profile (left) and pathway enrichment analysis (right) of γδ T cells (cluster 15) from the colon and small intestine of CONV-R and GF WT and PolgA^mut^ mice. **(E)** Gene expression profile (left) and pathway enrichment analysis (right) of macrophages (cluster 8) from the colon and small intestine of CONV-R and GF WT and PolgA^mut^ mice. Data shown as mean ±SEM. Statistical significance was determined using two-way ANOVA with Tukey’s correction for multiple comparisons; *p < 0.05, **p < 0.01, ***p < 0.001.

To further dissect myeloid cells transcriptional remodeling by microbiota and mitochondrial dysfunction, scRNAseq clusters 6, 7, 8, 11, and 24 were subsetted and re-analyzed at higher resolution (Figure S4D). This in-depth profiling revealed a marked decrease in tissue-resident macrophages (TRMs; cluster4) in PolgA^mut^ CONV-R mice, accompanied by a concomitant expansion of recently infiltrated inflammatory monocytes (cluster 3) (Figure S4D-F). This reciprocal shift suggests inflammatory reconstitution of the macrophage compartment, potentially driven by destabilization of the resident niche and compensatory recruitment of pro-inflammatory monocyte-derived cells. Together, these data indicate that microbiota-dependent signals promote expansion and activation of recently infiltrated inflammatory monocytes in PolgA^mut^ mice. Finally, we set out to evaluate whether GF conditions could establish a long-term functional imprinting on immune cells. We differentiated bone marrow-derived macrophages (BMDM) from both CONV-R and GF WT and PolgA^mut^ mice in classically (M1) and alternatively activated (M2) macrophages. Expression of key functional markers in both populations was comparable between CONV-R and GF conditions (Figure S4G), suggesting that genetic factors may play a more prominent role in shaping function when direct interactions with the microbiota are absent. Overall, the absence of microbiota improves the phenotype of PolgA^mut^ mice not only by decreasing the frequencies of immune cell infiltrating the intestinal tissue but by fundamentally reshaping their transcriptional profile diminishing their inflammatory potential.

## DISCUSSION

Microbiota dysbiosis has recently been recognized as a hallmark of aging ^7,8^. Several studies performing fecal matter transplantations from young to old WT ^12,29^ or progeric ^13^ animals have shown a positive impact on aging measurements, suggesting that microbiota dysbiosis is a contributing factor towards aging through regulation of inflammatory and immune responses. Yet, the opposite can also occur, where the transplantation of old microbiota to young mice increases neurogenesis and pro-longevity signaling^30^. In our work, we observed a significant interaction of genotype x microbiota composition over time, suggesting that mitochondrial dysfunction directly influences gut microbial community structure with age. Our findings are consistent with previous studies in PolgA^mut^ mice also reporting an expansion of Firmicutes, in particular *Ligilactobacillus murinis* and *Lactobacillus reuterii* species, at the detriment of other taxa ^31^. Of note, *Ligilactobacillus*, a genus associated with improved intestinal immunity, barrier function and metabolic homeostasis ^32^, was the most important genus for age prediction in a study of 2997 metagenomes collected longitudinally from 913 deeply phenotyped, genetically diverse mice ^4^. Altogether, these studies suggest that the accumulation of *Ligilactobacillus* may be a microbial biomarker of both normal and accelerated aging in mice. These parallels between WT and progeric mice support the selection theory, suggesting that aging and host mitochondrial dysfunction actively shape microbiota drift, rather than occurring solely by stochastic environmental exposures during aging as proposed by the neutral theory ^4^. Finally, irrespective of age-related dysbiosis or the enrichment of health-associated taxa, our data crucially indicate that the presence of a microbiota exerts an overall direct negative causal effect on lifespan.

According to a central tenet of modern biogerontology, aging is recognized as the consequence of biological programs that optimize early-life fitness at the expense of late-life maintenance. The antagonistic pleiotropy (AP) theory of aging postulates that traits conferring developmental, reproductive or survival advantages in youth are retained by natural selection, even if they become detrimental later in life ^33^. Within this framework, mechanisms that promote rapid growth, efficient nutrient extraction, and robust immune responsiveness early on may progressively drive metabolic imbalance and tissue decline with age. Here, we propose that the impact of gut microbiota on host physiology represents a case of AP. For example, gut microbiota promote postnatal growth in undernourished mice by supporting IGF-1, and insulin signaling through bacterial cell wall components acting via NOD2 in intestinal epithelial cells ^34^. Likewise, the capacity for reproduction in GF mice remains inferior to their conventional counterparts despite improvement of feeding or rearing conditions ^35^, with the microbiota extending the reproductive lifespan of mice by safeguarding the ovarian reserve ^36^. Our data and studies from GF invertebrates and mammals convincingly show that the absence of microbiota in WT animals robustly extends lifespan in an evolutionarily conserved manner^12,17,37–39^. Furthermore, GF status was recently shown to induce premature aging-like molecular signatures in young mouse tissues, but delay aging features in old mice across multiple cell types ^40^, consistent with our physiological and phenotypic observations. Altogether, these studies imply that while the role of microbiota during early life is beneficial, its relationship with their hosts is also physiologically costly and contributes to shortening lifespan.

So why is this symbiotic relationship costly with age? Our data highlight that the presence of a microbiota intensifies immune proteome remodelling in the small intestine and colon, promoting inflammatory myeloid reprogramming, plus significantly decreasing respiratory chain subunit expression systemically while also activating mTOR signaling in peripheral tissues such as muscle. Thus, the absence of microbiota positively impacts three hallmarks of aging – dysfunctional mitochondria, deregulated nutrient signaling and inflammaging. Of note, we observe that GF status improves mitochondrial bioenergetics through increased oxidative capacity and aerobic metabolism, shown by higher levels of SDH activity in both WT and PolgA^mut^ animals and by rescued expression of several respiratory chain subunits. Furthermore, the absence of microbiota inhibits mTOR activation and decreases translation. A similar mechanism was observed in flies where *L. plantarum* exerts its beneficial effect on systemic growth upstream of mTOR through a BCAA-dependent mechanism ^41^. We find that BCAA degradation is significantly and uniquely enriched in our data, being upregulated in several organs of GF PolgA^mut^ animals compared to CONV-R. This is possibly a compensatory mechanism to obtain propionyl-CoA, an important mitochondrial substrate through conversion to acetyl-CoA or succinyl-CoA that would normally be derived from the gut microbiota production of the SCFA propionate (Figure S1B,C). Remarkably, an epistatic interaction is observed between dietary restriction (DR) and GF status, where both interventions alone extend lifespan but the combination does not, or even shortens lifespan in both mice and rats ^38,39^. Such epistatic effects suggesting common mechanistic overlap between seemingly distinct interventions have been explained by their role on maturation of the immune system ^39^. DR was shown to delay age-related decline in immune function with higher immunological activity in old age compared to fully fed mice. Recently, the gut commensal *P. goldsteinii* was shown to drive age-associated cognitive decline through the production of medium-chain fatty acids causing peripheral myeloid cell inflammation ^42^. Likewise, our data further confirm that the metabolic reprogramming of the immune system by the microbiota also plays a role in modulating organismal lifespan.

Previous association studies in which gut microbial features correlated with multiple health parameters such as body composition, immune status and frailty, showed no clear association with lifespan itself ^4^. A possible interpretation is that the physiological cost of maintaining a microbiota is intrinsically integrated into host life history, such that microbial contributions to host fitness are offset by this baseline burden. In this framework and in the context of our data, microbiota-targeted interventions may alleviate aspects of this cost and improve healthspan, but are unlikely to extend lifespan beyond the constraints imposed by host-microbe coexistence.

## SUPPLEMENTAL FIGURE LEGENDS

**Figure S1.**
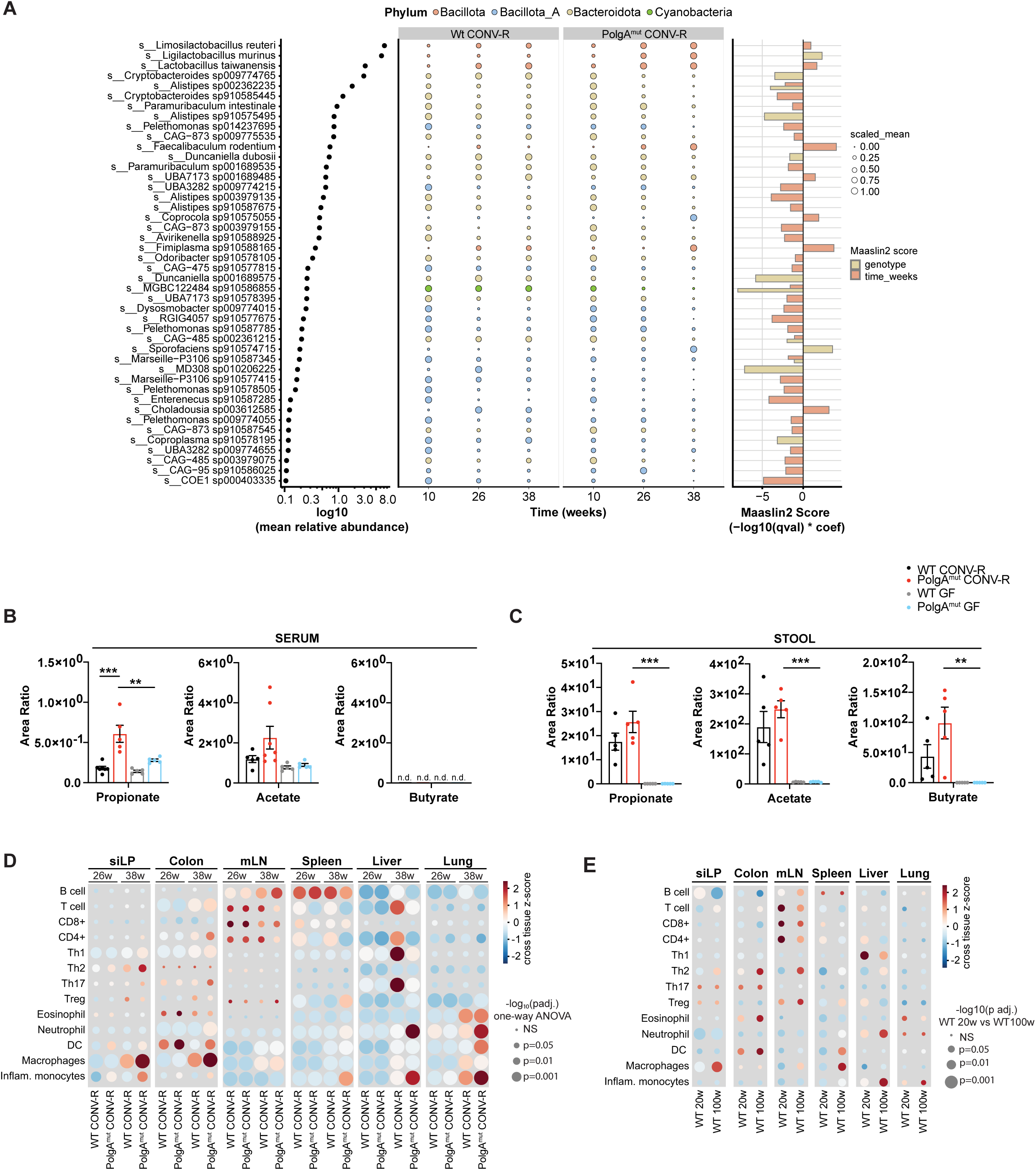
Related for Figure 1. Metagenomic profiles during aging and in a mitochondrial deficient mouse model. **(A)** Genotype- and age-associated changes in metagenome assemble genome (MAG) abundance. **(B)** SCFA quantification in serum collected from 38-week-old CONV-R and GF WT and PolgA^mut^ mice. **(C)** Propionate, acetate and butyrate quantification in stool collected from 26-week-old CONV-R and GF WTand PolgA^mut^ mice. **(D)** Z-score–normalization across tissues frequencies of tissue-infiltrating immune cells in the indicated organs of 26- and 38-week-old CONV-R WT and PolgA^mut^ mice. **(E)** Z-score–normalized normalization across tissues frequencies of tissue-infiltrating immune cells in the indicated organs of 20- and 100-week-old CONV-R WT mice.

**Figure S2.**
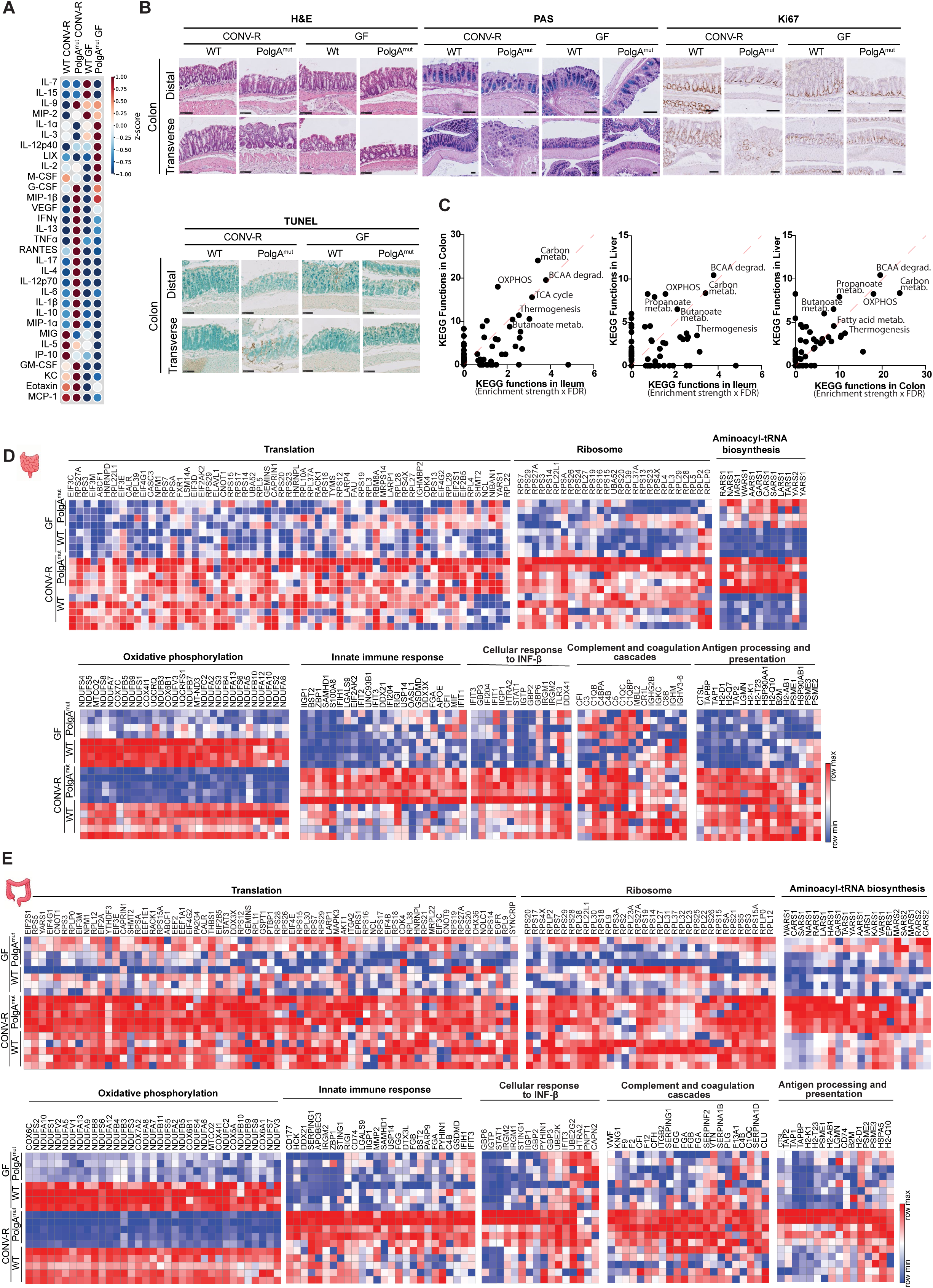
Related for Figure 2. Multi-organ proteomic and inflammatory landscape of PolgA^mut^ mice. **(A)** Z-score-normalized serum concentration levels of the indicated cytokines in 38-week-old CONV-R and GF WT and PolgA^mut^ mice. **(B)** Representative images of transverse and distal colon sections from 38-week-old CONV-R and GF WT and PolgA^mut^ mice stained with H&E, PAS and Ki67. **(C)** Tissue pairwise interaction of KEGG functional categories. **(D)** Heatmaps depicting levels of protein involved in indicated pathways in small intestine samples from 38-week-old CONV-R and GF WT and PolgA^mut^ mice. **(E)** Heatmaps depicting levels of protein involved in indicated pathways in colon samples from 38-week-old CONV-R and GF WT and PolgA^mut^ mice.

**Figure S3.**
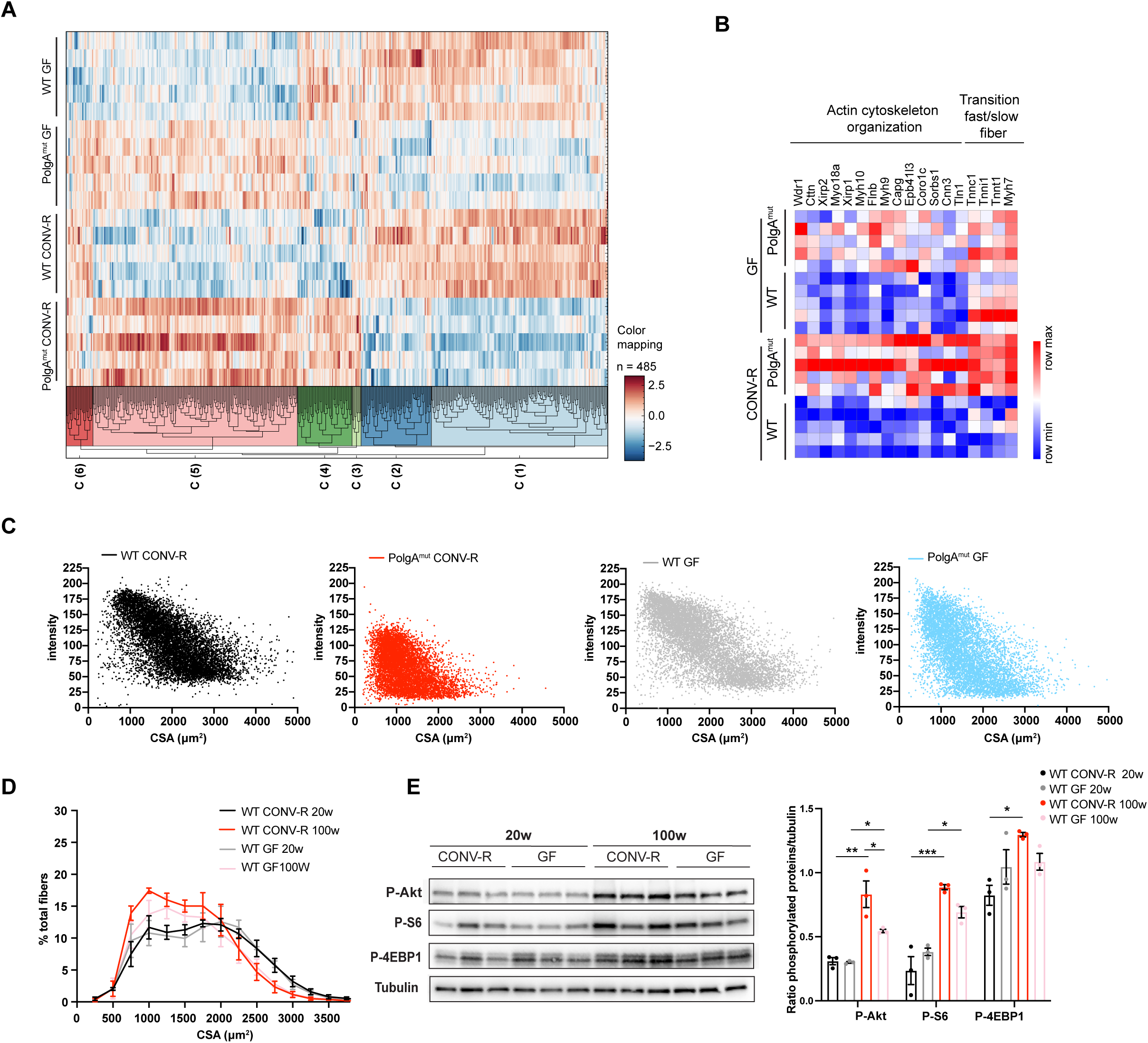
Related for Figure 3. Muscle proteomics and metabolic profile in PolgA^mut^ mice. **(A)** Subset of genes after two-way ANOVA comparing CONV-R and GF WT and PolgA^mut^ mice. **(B)** Heatmap of proteins associated with actin cytoskeleton organization and fast-to-slow muscle fiber transition across experimental groups. **(C)** Correlation analysis between muscle fiber cross-sectional area (CSA) and SDH staining intensity in the indicated experimental groups. **(D)** Frequency distribution of muscle fiber CSA (μm^2^) in CONV-R and GF WT mice at 20 and 100 weeks of age. **(E)** Total protein extracts from muscle of CONV-R and GF WT mice at 20 and 100 weeks of age, immunoblotted with the indicated antibodies. Data are presented as mean ±SEM. Statistical significance was determined using two-way ANOVA with Tukey’s correction for multiple comparisons; *p < 0.05, **p < 0.01, ***p < 0.001.

**Figure S4.**
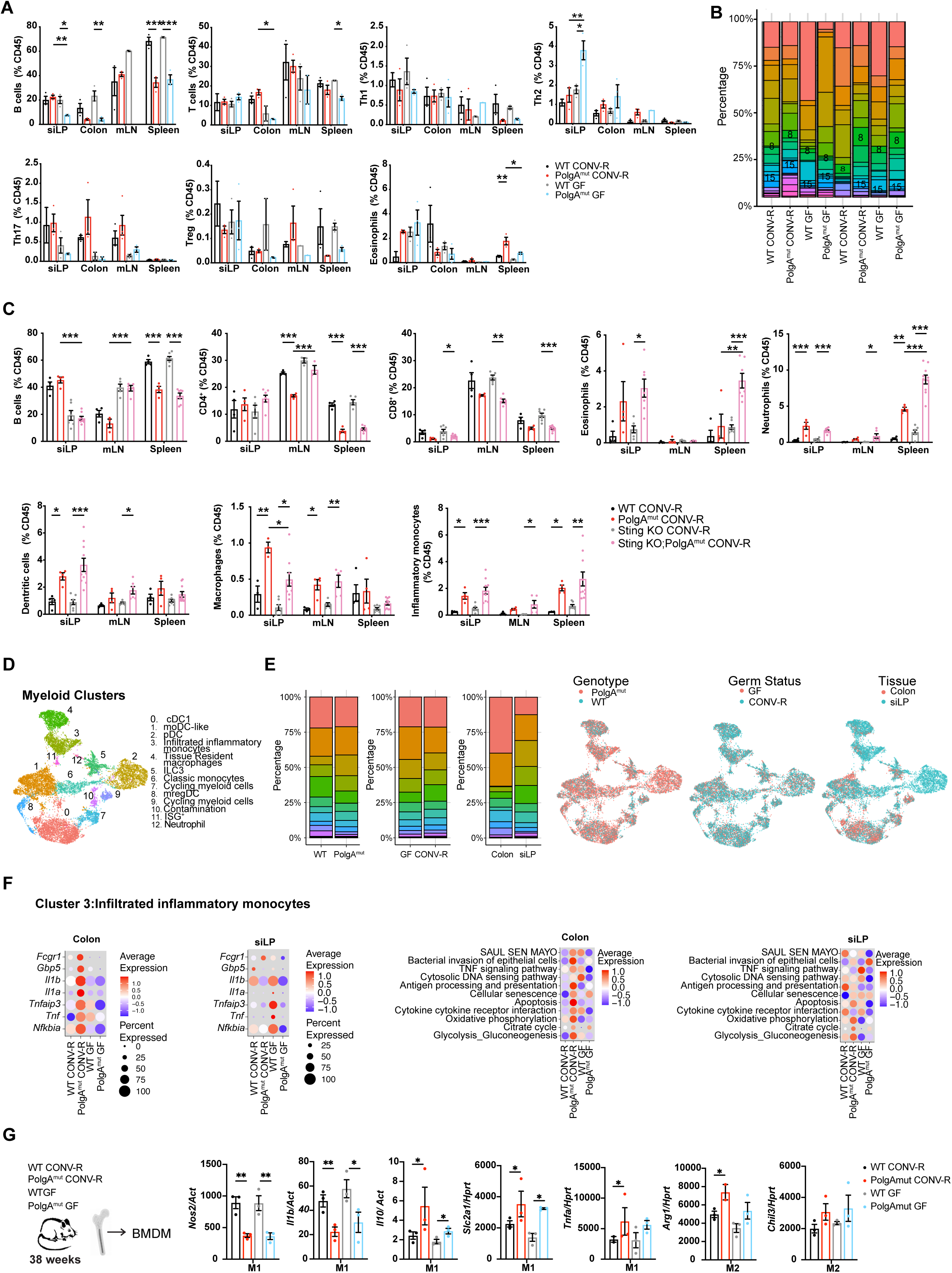
Related for Figure 4. scRNAseq and cytometry analysis of gut immune infiltrate in PolgA^mut^ mice. **(A)** Multi-organ immunophenotyping of 38-week-old CONV-R and GF WT and PolgA^mut^ mice with frequencies of tissue-infiltrating immune cells. **(B)** Frequencies of annotated cell clusters in CD45^+^ cells sorted from the colon and small intestine of CONV-R and GF WT and PolgA^mut^ mice. **(C)** Multi-organ immunophenotyping of 38-week-old CONV-R WT, PolgA^mut^, STING KO and STING KO;PolgA^mut^ mice with frequencies of tissue-infiltrating immune cells. **(D)** UMAP visualization of the myeloid cell clusters distribution after subset and reanalysis at higher resolution and cell population annotation. **(E)** Frequencies of annotated cell clusters in myeloid cells in the colon and small intestine of CONV-R and GF WT and PolgA^mut^ mice (left). UMAP visualization distribution per genotype, germ status and tissue (right). **(F)** Pathway enrichment analysis of transcriptional profile of recently infiltrated inflammatory monocytes (cluster 3) from the small intestine and colon of CONV-R and GF WT and PolgA^mut^ mice. **(G)** Analysis of bone marrow-derived macrophages (BMDMs) cultured from CONV-R and GF WT and PolgA^mut^ mice. Data shown as mean ±SEM. Statistical significance was determined using two-way ANOVA with Tukey’s correction for multiple comparisons; *p < 0.05, **p < 0.01, ***p < 0.001.

## ACKNOWLEDGEMENTS

FC was supported by the Wellcome Trust/Royal Society (102532/Z/12/Z and 102531/Z/13/A), the DFG German Research Foundation (EXC 2030 -390661388) and by i-HEAD (PB-NRW-2022-ANTRAG-025). FC, MC and FS were supported by Center for Molecular Medicine Cologne (IPFP-B02). MC was supported by the DFG German Research Foundation (269925409-B12) and Center for Molecular Medicine Cologne (IPFP-B03 and CAP21). FS was supported by the Deutsche Forschungsgemeinschaft (DFG, German Research Foundation) SFB-1403–414786233. DMM was supported by the Leverhulme Trust (RPG-2022-299). This work was also supported by the large instrument grant INST 1856/71-1 FUGG by the German Research Foundation (DFG Großgeräteantrag). We acknowledge computational resources and support provided by the Imperial College Research Computing Service (http://doi.org/10.14469/hpc/2232). We thank Christoph Kaleta and Helena Cochemé for critically reading the manuscript and for their insightful comments and suggestions.

## AUTHOR CONTRIBUTIONS

FS and FC conceptualized the research. FS performed experimental procedures and analysed data. DMM, JW contributed to metagenomic computational analysis. SB performed the metabolomic identification and analysis. NGZT and GGK and MP supported histological analysis. AR, AS, JVL, KM, contributed to experimental procedures. CDN, LN and BB performed ex-vivo muscle measurements. SL and MK performed proteomics identification and analysis. DES supported with ScRNAseq analysis. FS and FC wrote the manuscript. FS, MC and FC participated in the interpretation of the main findings. FS and FC supervised the research. FS, MC and FC secured funding. All authors read, provided feedback and approved the final manuscript.

## COMPETING INTERESTS

The authors declare no competing interests.

## MATERIAL AND METHODS

### Mice

PolgA^mut^ and Sting KO;PolgA^mut^ were kindly donated by Dr. Aleksandra Trifunovic. Germ-free (GF) mice were generated by Taconic Biosciences via *in vitro* fertilization. Embryos were transferred into GF Swiss Webster pseudopregnant recipient females, which were subsequently housed in germ-free isolators. Recipient females then delivered germ-free offspring. GF mice were maintained in class biologically clean (CBC) isolators supplied with HEPA-filtered air and a controlled, stratified pressure system. Animals were housed in an open cage system within the isolators. Food, water, and bedding were sterilized prior to use, and animals were provided food and water *ad libitum*. Sterilization procedures were performed by autoclaving or by transfer through a hydrogen peroxide (H₂O₂) fumigation airlock. Mice were maintained at the ivRF CECAD Research Institute (Cologne, Germany) under specific-pathogen-free (SPF) conditions or GF conditions and cared for according to the Institutional Animal Use and Care Guidelines and under a 12-h dark–12-h light cycle. SPF mice were mantained in individually ventilated cages (Greenline GM500, Tecniplast) at 22 ±2°C and a relative humidity of 55 ±5%. All mice had unlimited access to water and fed a standard chow diet (Sniff, #V1554-300 or #V1185-300) *ad libitum*. Health monitoring program followed FELASA recommendations ^43^. Briefly, germ-free NMRI mice are used as bedding sentinels receiving dirty bedding from up to 100 other cages weekly for three months before being tested for all pathogens and opportunists recommended in the guidelines. Breeding of the animals was approved by Landesamt für Natur, Umwelt und Verbraucherschutz Nordrhein-Westfalen (LANUV NRW).

### Isolation of lymphocytes from the small intestine

Isolation of cells from the small intestine was performed as follows. Upon harvesting, fat tissue bordering the small intestine and Peyer’s patches were removed. Small intestines were cut open, washed with ice-cold PBS 1X + 25 mM HEPES (Sigma-Aldrich, #83264), cut into 2 cm-long pieces and collected in medium containing RPMI1640 (Thermo Fisher Scientific, #21875158), 25 mM HEPES (Sigma-Aldrich, #83264), 3% FCS (Gibco, #26140079), 4 mM glutamine (Life Technologies, #25030-024), 1% penicillin/streptomycin (Life Technologies, #15140-122) and 55 μM β-mercaptoethanol (Thermo Fisher Scientific, #21985023) (hereafter named 3% FCS medium). Small intestines were incubated in 3% FCS medium supplemented with 5 mM EDTA (Thermo Fisher Scientific, #15575020) and 1 mM dithiothreitol (DTT) (Sigma Aldrich, #10197777001) for 25 min, at 37°C, 5% CO_2_ and agitation (1^st^ incubation). After the incubation, the leftover small intestine pieces were thoroughly (4 times) washed by shaking with ice-cold medium containing RPMI1640, 25 mM HEPES (Sigma-Aldrich, #83264), 4 mM glutamine, 1% penicillin/streptomycin and 55 μM β-mercaptoethanol (hereafter named serum-free medium) supplemented with 2 mM EDTA. Washed small intestine pieces were then finely chopped and incubated in serum-free medium added with 100 μg/mL Liberase TL (Roche, #05401020001) and 50 μg/mL DNAse I (Sigma-Aldrich, #10104159001) for 35 min, at 37°C, 5% CO_2_ and agitation (2^nd^ incubation). After digestion, suspensions were strained through 70 μm strainers, washed and kept as lamina propria (LP) fraction. LP fractions were enriched for leukocytes using a three-layer Percoll (Merck, #17089101) gradient (Percoll 75%, 40% and 30%) centrifuged for 20 min, at room temperature, 680 x *g*, without any centrifuge acceleration and brake. The leukocyte layer between Percoll 75% and 40% was collected as enriched LP fraction, extensively washed and suspensions used for downstream applications.

### Isolation of lymphocytes from the colon

Cell suspensions from the colonic lamina propria were prepared as follows: briefly, colons were washed twice in RPMI 5% FCS supplemented with 5 mM EDTA and then once in RPMI 5% FCS containing 15 mM HEPES in a shaking incubator. Each wash was performed at 200 rpm 20 min at 37°C. Digestion of the colon tissue was performed in RPMI 5% FCS containing 1 mg/mL type VIII collagenase (Sigma Aldrich, #C2139) and 40 mg/mL DNase I (Sigma-Aldrich, #10104159001) at 37°C for 60 min on a shaker at 200rrpm. Supernatants were then filtered on a 70μm strainer and a three-layered discontinuous Percoll gradient (Percoll 75%, 40% and 30%) was used to obtain an enriched leukocyte fraction (layer between Percoll 75% and 40%).

### Isolation of lymphocytes from the liver and lung

Whole liver was collected in ice-cold complete medium, finely chopped and incubated in RPMI1640 supplemented with 1 mg/mL collagenase IV (Sigma-Aldrich, #C2139) and 50 μg/mL DNAse I (Sigma-Aldrich, #10104159001) for 45 min, at 37°C, 5% CO_2_ and agitation. After digestion, suspensions were strained through 70 μm strainers, washed and enriched for leukocytes using Percoll 33% and centrifuging them for 20 min, at room temperature, 680 x *g*. The leukocyte pellet was collected, extensively washed, red blood cell lysis was performed and suspensions used for downstream applications. Lung digestions were performed as follows. Lungs were collected in ice-cold complete medium, finely chopped and incubated in complete medium added with 2 mg/mL collagenase IV (Sigma-Aldrich, #C2139) and 50 μg/ml DNAse I (Sigma-Aldrich, #10104159001) for 45 min, at 37°C, 5% CO_2_ and agitation. After digestion, suspensions were strained through 70 μm strainers, washed and enriched for leukocytes using a three-layer Percoll gradient (Percoll 75%, 40% and 30%) centrifuged for 20 min, at room temperature, 680 x *g*, without any centrifuge acceleration or brake. The leukocyte layer between Percoll 75% and 40% was collected, extensively washed and suspensions used for downstream applications.

### Isolation of lymphocytes from spleens and mesenteric lymph node (mLN)

Spleens and lymph nodes were harvested and mashed with a syringe plunger in a petri dish with 5 mL of T cell medium (TCM; 1640 RPMI with 10% fetal calf serum, 4 mM L-glutamine, 1% penicillin/streptomycin, 55 μM β-mercaptoethanol). The cell suspension was filtered through a 70 μm strainer, centrifuged at 400 x *g* for 5 min at 4°C and resuspended in 1 mL of LCK (Thermo Fisher Scientific, #A1049201) to lyse red blood cells. After 3 min TCM was added, and the cell suspension was centrifuged at 400 x *g* for 5 min at 4°C. The pellet was resuspended in 1 mL of TCM and filtered through 40 μm cell strainers.

### Bone marrow derived macrophages (BMDMs) culture

Mature bone marrow derived macrophages (BMDMs) were obtained at day 7 of culture of bone marrow cells in 20 ng/mL CSF-1 (Peprotech) in RPMI medium containing 10% fetal bovine serum, 4 mM L-glutamine, 100 U/mL penicillin/streptomycin (complete RPMI). BMDMs were stimulated in complete RPMI supplemented with 20 ng/mL CSF-1 (Peprotech) for additional 24 h with a combination of 20 ng/mL IL-4 (Peprotech) for M2 differentiation or 50 ng/mL IFN-γ (Peprotech) and 20 ng/mL LPS (SIGMA) for M1 differentiation under 5% CO_2,_ atmospheric oxygen, at 37°C in a humidified incubator.

### Flow cytometry

Fluorochrome-conjugated antibodies were purchased from eBioscience, BD Bioscience, BioLegend. Cells were detected using Fortessa flow cytometers and analysed using FlowJo software (FlowJo). Cell viability was quantified by flow cytometry using LIVE/DEAD^TM^ aqua or near-IR dyes (Invitrogen) following manufacturer instructions. Cell discrimination was done applying the following gating strategy: 1) lymphoid cells were identified based on forward scatter (FSC) and side scatter (SSC) parameters; 2) single cells were discriminating plotting the area of the FSC vs the width of FSC; 3) live/dead staining was uses to exclude dead cells from analysis: live cells are negative for the live/dead staining; 4) the different cell populations were identified based on the surface markers expression.

### RT-PCR

RNA was isolated using a RNeasy Mini Kit (#74106, Qiagen) according to manufacturer’s instructions. RNA concentration was measured using Nanodrop 2000 (Thermo Fisher). cDNA synthesis was performed using High-Capacity cDNA Reverse Transcription Kit (#4368813, Thermo Fisher) and RNasin Plus RNase Inhibitor (#N2615, Promega), according to manufacturer’s instructions. Per reaction, 450-1000 ng RNA was used. RT-qPCR was performed in 384-wells plates using iTaqMan Universal Probes SuperMix (#1725134, Bio-Rad), 10 ng cDNA per sample and 20X TaqMan Gene Expression Assay (Nos2: Mm00440502_m1; Il-1β: Mm00434228_m1; Il-10: Mm01288386_m1; Slc2a1: Mm00441480_m1; Tnfα: Mm00443258_m1; Arg1: Mm00475988_m1; Chil3: Mm00657889_mH; Hprt: Mm03024075_m1; Thermo Fisher). RT-qPCRs were performed with a QuantStudioTM 5 Real-Time PCR 384 well system (#A28140, Thermo Fisher). Gene expression was normalized to Hprt expression. Fold change was calculated according to the delta-Ct method (2^-dCt^) or delta-delta Ct method (2^-ddCt^).

### scRNAseq

CD45^+^ single cells were sorted with BD FACSAria Cell Sorter and prepared for single cell RNA sequencing using the 10X Genomics 3’scRNA v3 system. Libraries were sequenced on NovaSeq 6000 at 250 M reads/sample. Samples were demultiplexed, quality checked, filtered and aligned genome build GRCm38 using pre-established pipelines implemented via cellranger v.8.0.1. Resulting raw read count matrix of barcodes corresponding to cells and features corresponding to detected genes were processed, analyzed and visualized in R v.4.3.1 using Seurat v.5^39^ with default parameters in all functions, unless specified. Poor quality cells, with low total unique molecular identifier (UMI) counts and high percent mitochondrial gene expression, were excluded. Filtered samples were normalized using a regularized negative binomial regression (SCTransform)^40^ and integrated with Harmony^41^, using 50 principal components. Integrated gene expression matrices were visualized with a UMAP^42^ as a dimensionality reduction approach. Resolution for cell clustering was determined by evaluating hierarchical clustering trees at a range of resolutions (0 - 1.2) with Clustree^43^ and by calculating a silhouette score, selecting a value inducing minimal cluster instability and maintaining a high proportion of cells with a positive silhouette score. Differentially expressed genes between clusters were identified as those expressed in at least 10% of cells with a greater than +1 log fold change and an adjusted p value of less than 0.01, by transforming read count matrices to pseudobulk using Libra (https://github.com/neurorestore/Libra) and then applying DESeq2^44^ v.1.36 with default parameters. Artificial replicates for differential gene expression analysis were generated by randomly partitioning single cell dataset into three tiles. Ribosomal protein genes were excluded from results. Cluster specific genes were explored for pathway enrichment using StringDB^45^. Gene set scores were calculated using the UCell with default parameters^46^. scRNAseq dataset is available with accession number GSE325681.

### Cytokine measurements

Cytokine concentration was measured from mouse using Luminex xMAP technology for multiplexed quantification of 32 Mouse cytokines, chemokines, and growth factors. The multiplexing analysis was performed using the Luminex™ 200 system (Luminex, Austin, TX, USA) by Eve Technologies Corp. (Calgary, Alberta). Thirty-two markers were simultaneously measured in the samples using Eve Technologies’ Mouse Cytokine 32-Plex Discovery Assay® (MilliporeSigma, Burlington, Massachusetts, USA) according to the manufacturer’s protocol. The 32-plex consisted of Eotaxin, G-CSF, GM-CSF, IFNγ, IL-1α, IL-1β, IL-2, IL-3, IL-4, IL-5, IL-6, IL-7, IL-9, IL-10, IL-12(p40), IL-12(p70), IL-13, IL-15, IL-17, IP-10, KC, LIF, LIX, MCP-1, M-CSF, MIG, MIP-1α, MIP-1β, MIP-2, RANTES, TNFα, and VEGF. Assay sensitivities of these markers range from 0.3 – 30.6 pg/mL for the 32-plex. Individual analyte sensitivity values are available in the MilliporeSigma MILLIPLEX® MAP protocol.

### Metagenomics

Raw reads from BGI DNBSEQ™ sequencing platform were demultiplexed and their quality evaluated with FastQC v.0.12 (https://www.bioinformatics.babraham.ac.uk/projects/fastqc) and multiqc v.1.14^44^. Sequences adapters were trimmed and filtered following standard procedures by using trimmomatic v.0.39^45^. Sequences coming from the host were mapped against the Mus musculus reference genome (GRCm38) and dropped using bowtie2 v.2.5.4 ^46^. After pre-processing the reads, taxonomy assignment was done with MetaPhlAn v.4.06 using by default parameters and the Oct22 database ^47^ to capture to the species level with high quality. Taxonomy and OTU table were then processed and analysed in R. The main analysis was done using tidyverse v.2.0.1 and microeco v.1.6.0 ^48^. To capture the beta-diversity of the samples, Jaccard distances were used to calculate the Principal Coordinate Analysis (PCoA). Statistical association of bacterial relative abundance with genotype and husbandry was done with Maaslin2 in R ^49^. Microbial temporal variability was calculated as in Marti et al. ^20^, fitting a power law where V (the main indicator of temporal variability) and beta were the parameters to fit. Taylor’s law parameters were weighted by applying a bootstrap (1000 iterations) where the mean (x-variable) was resampled by applying random Gaussian with error equal to the SEM of the measurement. Graphical representation of the results was done with the R package ggplot2 v.3.5.1.

### Analysis of short-chain fatty acids (SCFAs)

Levels of acetate, propionate and butyrate in mouse stool and serum were determined by Anion-Exchange Chromatography coupled to Electrospray Ionization High-Resolution Mass Spectrometry (IC-ESI-HRMS): Approximately 30 mg of snap-frozen mouse stool were homogenized in ice-cold methanol/water 4:1 (v/v/v) (1 mg/20 µL) using the Precellys 24 Homogenisator (Peqlab) at 6,400 rpm for two times 10 s with a 5-s pause. To 360 µL stool homogenate, 20 µL of a mixture of isotope-labeled internal standards in Milli-Q water (100 µM sodium acetate-^13^C_2,_d_3;_ 100 µM propionic acid-d_6_; 100 µM butyric acid-d_7,_ Sigma-Aldrich) were added. To 100 µL serum, 400 µL ice-cold methanol and 20 µL of the internal standard mixture were added. The samples were vortexed for 20 sec and incubated at -70°C for 30 min. The samples were then extracted using a pre-cooled ThermoMixer (Eppendorf) at 4°C and at 3,000 rpm for 15 min. After centrifugation (16,100 RCF, 20 min, 4°C), the supernatants were removed, transferred to new tubes, vortexed for 20 sec and again incubated at -70°C for 30 min. After centrifugation (16,100 RCF, 20 min, 4°C), the supernatants were removed, transferred to new tubes and dried under a stream of nitrogen. The dried extracts were resolved in 100 µL of Milli-Q water. After mixing and centrifugation (16,100 RCF, 20 min, 4°C), 80 µL of the supernatants were transferred to autoinjector vials and immediately measured.

IC-HRMS analysis was performed using a Dionex Integrion RFIC system (Thermo Scientific) equipped with a Dionex IonPac AS11-HC column (2 mm × 250 mm, 4 μm particle size, Thermo Scientific) and a Dionex IonPac AG11-HC guard column (2 mm × 50 mm, 4 μm, Thermo Scientific) and coupled to a Q Exactive HF quadrupole-orbitrap mass spectrometer (Thermo Scientific). 5 µL of sample were injected using a Dionex AS-AP at 5°C. The IC was operated at a flow rate of 0.38 mL/min with a potassium hydroxide gradient which was produced by an eluent generator with a potassium hydroxide cartridge and Milli-Q water. The gradient started with 10 mM KOH over 3 min, 10−50 mM from 3 to 12 min, 50−100 mM from 12 to 19 min, held at 100 mM from 19 to 25 min, and re-equilibrated at 10 mM for 3 min. An Dionex ADRS 600, 2 mm suppressor was operated with 95 mA, and methanol/water 95:5 (v/v) was used to produce a make-up flow at a flow rate of 0.15 ml/min ^50^. The mass spectrometer was operated in the negative ion mode. Full MS scans in the range of m/z 50-120 were acquired with a resolution of 15,000, an Automatic Gain Control (AGC) target value of 3 x 10^6^ and a maximum injection time (IT) of 240 ms. Spectrum data were collected in the profile mode. The ESI source was operated with flow rates for sheath gas, auxiliary gas, and sweep gas of 30, 14 and 3, respectively. The spray voltage setting was 2.75 kV, the capillary temperature 230°C, the S-lens RF level 60, and the auxiliary gas heater temperature 380°C. The exact m/z traces of acetate and propionate and the isotope-labeled internal standards were extracted and integrated using the TraceFinder v.5.1 software (Thermo Scientific). Endogenous SCFAs were quantified by normalizing their peak areas to those of the internal standards.

### Proteomics

Tissue powder generated under liquid nitrogen were lysed in 8 M urea in 50 mM triethylammonium bicarbonate (TEAB) supplemented with protease inhibitor cocktail (Roche). Chromatin was disrupted by sonication (Bioruptor, Diagenode; 10 min, 30 s on/off cycles) or alternatively by treatment with Benzonase HC nuclease (25 U per 5 x 10^5^ cells; Merck). Lysates were cleared by centrifugation (20,000 × *g*, 15 min), and protein concentrations were determined using a standard protein assay.

For digestion, 50 µg of protein per sample were reduced with 5 mM dithiothreitol (DTT; Sigma-Aldrich) for 1 h at 25°C and alkylated with 40 mM chloroacetamide (CAA; Sigma-Aldrich) for 30 min in the dark. Where indicated, proteins were pre-digested with Lysyl Endopeptidase (Lys-C; Wako, #125-05061) at an enzyme-to-substrate ratio of 1:75 for 4 h at 25°C. Samples were subsequently diluted with 50 mM TEAB to decrease the urea concentration to ≤2 M and digested overnight with sequencing-grade trypsin (enzyme-to-substrate ratio 1:75; Promega) at 25°C. Digestion was stopped by acidification with formic acid to a final concentration of 1%. Peptides were purified using SDB-RP StageTips prior to LC-MS/MS analysis. The mass spectrometry proteomics data have been deposited to the ProteomeXchange Consortium via the PRIDE partner repository with the dataset identifier PXD075317.

Samples were analyzed in DIA-NN 1.8.1^51^. A Swissprot mouse canonical database (UP589, downloaded 18/06/2020) was used for library building with settings matching acquisition parameters and the match-between-runs function enabled. Here, samples are directly used to refine the library for a second search of the sample data. DIA-NN was run with the additional command line prompts “—report-lib-info” and “—relaxed-prot-inf”. Further output settings were: filtered at 0.01 FDR, N-terminal methionine excision enabled, maximum number of missed cleavages set to 1, minimum peptide length set to 7, maximum peptide length set to 30, mininimum precursor m/z set to 400, maximum precursor m/z set to 1000, cysteine carbamidomethylation enabled as a fixed modification. Afterwards, DIA-NN output was further filtered on library q-value and global q-value <= 0.01 and at least two unique peptides per protein using R (v.4.1.3). Finally, LFQ values calculated using the DIA-NN R-package. Afterwards, analysis of results was performed in Perseus v.1.6.15 ^52^.

### Proteomics of muscle tissue

Tissue samples were resuspended in 4% SDS in PBS, homogenized by heating for 10 min at 95°C, and sonicated with a Bioruptor sonicator; the sonication settings were 21°C water with 10 cycles of 30s on and 30s off. Protein concentrations were determined using the Pierce Protein Assay (Thermo Scientific), then 250 µg protein extract was reduced and alkylated for 10 min at 70°C using 5 mM TCEP and 15 mM CAA. Protein digestion was performed following the SP3 protocol ^53^. In brief, 250 µg of both hydrophylic and hydrophobic beads were added to the sample and bound by adding 1:1 volume of acetonitrile (ACN). After incubation for 8 min, magnetic beads were immobilized and washed 2x with 70% ethanol and ACN. Proteins were digested with trypsin (substrate:enzyme ratio 100:1) overnight at room temperature. Peptides were acidified with TFA (pH < 2) and desalted using Sep-Pak C18 1 cc Vac Cartidge, 50 mg (Waters, Milford, US). Eluate was dried using a SpeedVec Concentrator Plus (Eppendorf). 4 µg peptides were used for proteomic analysis. In brief, 10:1 (w:w) Ti-IMAC were added Purified peptides were analysed were analyzed on an Orbitrap Eclipse Tribid Mass Spectrometer (Thermo) coupled with an easy nLC 1200 (Thermo). An in-house packed C18 analytical column (30 cm, 150 μm inside diameter, and 1.9-μm ReproSil-Pur C18 beads; Dr. Maisch, Germany) was used with an integrated column oven (50°C; PRSO-V1, Biberach, Germany). Phosphorylated peptides were analysed using a 90 min gradient (0-72 min: 4-30 %B, 73-80 min: 30-55 %B, 81-90 min: 55-95 %B). Four full MS scan was set to 120,000 at mass/charge ratio 390 to 1010 m/z, with AGC target at 300% and maximum injection time of 52 ms. DIA scans were performed from a precursor mass range m/z 400-550, 550-700, 700-850, 850-1000 with a resolution of 50,000. Proteome spectra were acquired within a 60 min gradient (0-48 min: 4-30 %B, 48-53 min: 30-55 %B, 53-6 min: 55-95 %B) and DIA using two full scans of 380-1050 with a resolution of 15K and two DIA scans (30×15 m/z) from a precursor mass range of 400-850 m/z with a resolution of 15L and a maximum injection time of 25 ms.

The mass spectrometry proteomics data have been deposited to the ProteomeXchange Consortium via the PRIDE partner repository with the dataset identifier PXD075562. Acquired spectra were processed using Spectronaut (v.19.1) using library free search against UniProt mouse database (April 2024). Data was further processed using R (v.4.2.2), with the libraries: tidyverse, data.table, magrittr, FactoMineR, factoextra and ggplot2, gprofiler, missForest, ggplot2. Data input was filtered for unique peptides, q-Value <0.01, Lib.Q.Value <0.01, PG.Q.Value < 0.01, Global.Q.Value < 0.01, Quantity.Quality > 0.7, Fragment.count >= 4. Statistical analysis was performed on the MaxLFQ normalised data with R (v.4.2.2), and Perseus (v.1.6.5.0) and visualized with Instant clue (v.0.10.10.20210315). Data completeness of 70% was calculated on each group. Missing values were imputed by random forest algorithm for groups with 70% data completeness, with more than 30% missing values random forest algorithm was downshifted of 0.3, width 1.5. ANOVA analysis and Welch’s T-test analysis was performed with an FDR with less than 0.05 and 500 randomizations. Quality control of iRT peptides were performed in Skyline-Daily (v.22.21.391).

### Single fiber tension measurement

Individual muscles were harvested and skinned as described in ^1^ while single fiber tension has been performed as reported in ^2^. In brief, a thin bundle of skinned fibers was dissected and incubated for 10 min in cold relax buffer (120 mM potassium propionate, 50 mM MOPS, 9 mM EGTA, 6 mM MgCl_2,_ 6 mM KH_2P_O_4,_ 5.2 mM ATP, 1 mM DTT, pH 7.0) containing 0.5% Triton X-100, then washed three times using fresh relaxing buffer. Single muscle fibers were mechanically separated using fine tweezers, clipped with aluminum T-clips, and mounted on an Aurora Scientific Permeabilized Fibers 802D setup (Aurora Scientific, Ontario, Canada). A thin flow of 5% toluidine blue and 8% glutaraldehyde was used to crosslink each end of the fiber^3^. After several washes with relaxing solution, the sarcomere length was adjusted to 2.5 μm using a camera mounted on an inverted microscope, and the diameter was measured. Fibers were activated at 7°C and moved to the following activating buffer at 17.5°C to develop maximal tension following the protocol named T-jump^4^. Relaxing, preactivating and activating buffers were adopted from^5^. Force was normalized to cross sectional area (CSA) using the measured diameter and assuming circular geometry.

### Tissue preparation for histology

Small intestines were dissected and washed with PBS. Small pieces (about 0.5 cm) were isolated proximal (after the stomach) and distal (before the caecum), the tissue was cut longitudinally and washed in PBS to remove feces. Intestinal tissue samples were rolled up from proximal to distal to form a Swiss roll and either fixed in 4% paraformaldehyde overnight at 4°C or embedded in TissueTek for cryo-sectioning.

### H&E staining of paraffin-fixed tissues

Paraffin-embedded 3-μm-thick intestinal tissue sections were deparaffinized with xylene and rehydrated with decreasing amounts of ethanol solutions (100%, 96% and 75%). Sections were stained for 2 min in haematoxylin, differentiated in tap water for 15 min, and incubated for 1 min in eosin. Stained sections were dehydrated using increasing amounts of ethanol solutions and fixed in xylene for 1 min. Slides were mounted with Entellan.

### PAS staining on paraffin-fixed tissues

Paraffin-embedded tissue sections were deparaffinized and rehydrated to distilled water. Sections were incubated with freshly prepared periodic acid solution for 5–8 min, followed by two washes in distilled water (5 min each). Slides were then incubated with Schiff’s reagent for up to 20 min and washed under running tap water for 10 min. Nuclei were counterstained with hematoxylin for 5 min and rinsed under running tap water for an additional 10 min. Sections were dehydrated through graded ethanol, cleared in xylol, and mounted using Eukitt or Entellan.

### Histology and immunohistochemistry

After samples from mice were fixed in 4% PFA, they were embedded in paraffin. Sections of 3 μm were subjected to histological analysis by haematoxylin and eosin (H&E) staining. To detect DNA fragmentation, TUNEL assay was performed by using a commercial kit (Abcam, ab206386) according to the manufacturer’s manual. For immunohistochemical analysis, slides were rehydrated and incubated with peroxidase blocking buffer (0.04 M sodium citrate, 0.121 M Na_2H_PO_2,_ 0.03 M NaN_3,_ 3% H_2_O_2_) for 10 min. Slides were washed and antigen retrieval was performed by digestion with proteinase K (10 μg/L) for 5 min in TEX-buffer (50 mM Tris, 1 mM EDTA, 0.5 % Triton X-100, pH 8.0) for CD45 and lysozyme or with sodium citrate buffer in a pressure cooker for 20 min for Ki67. Sections were washed with TBS supplemented with 0.1% Tween-20 (v/v) (TBS-T) and blocked for 60 min using commercial blocking buffer (ThermoFisher, #37536) supplemented with biotin from (1:100 dilution) from Avidin/Biotin blocking kit (Vector, #SP-2001) to block endogenous avidin. After sections were briefly washed with TBS-T, they were incubated with the primary antibody CD45 (eBioscience, #14-0451), lysozyme (Dako, #F0372) and avidin (1:100 dilution) from Avidin/Biotin blocking kit (Vector, # SP-2001) overnight at 4°C. A biotinylated secondary anti-rat antibody (Invitrogen, # D20697) or anti-rabbit (Invitrogen, #B2770) was incubated on the sections for 60 min. Stainings were visualized using the ABC Vectastain Elite Kit (Vector Laboratories) and DAB substrate (Abcam, #ab64238). Incubation times with DAB substrate were equal for all samples within one staining. Sections were then counterstained with haematoxylin for staining of the nuclei, dehydrated and mounted with Entellan. Histological sections were scanned using a Nanozoomer S360 (Hamamatsu) slide scanner. Images were analysed and processed using the Omero software package (https://openmicroscopy.org) and NDP.view2 Viewing software (Hamamatsu).

### Muscle histology, SDH staining and CSA

Histology was performed on 10-μm–thick cryosections: For hematoxylin and eosin staining, a hematoxylin solution according to Mayer (Sigma-Aldrich, S1275) and eosin purchased from Leica were used; succinate dehydrogenase (SDH) staining was obtained by incubating slices for 30 min in nitrotetrazolium blue chloride solution (N6876-500MG).CSA analyses were performed with SMASH (a MATLAB application) using a WGA staining for myofiber segmentation and myofiber contouring.

### Western Blot

Muscles were lysed in a buffer containing 50 mM Tris pH 7.5, 150 mM NaCl, 10 mM MgCl2, 0.5 mM DTT, 1 mM EDTA, 10% glycerol, 2% SDS, 1% Triton X-100, Roche Complete Protease Inhibitor Cocktail, and Roche Phospho-Stop Phosphatase Inhibitor Cocktail. Lysates were incubated at 70°C for 10 min and centrifuged at 16,000 x *g* for 15 min at 4°C. Cleared protein lysate were quantified using a BCA protein assay kit (Pierce #23225). Lysates were prepared in 2× Laemmli buffer with DTT and equal amounts of protein were resolved on 4–20% TGX gels (Bio-Rad). Proteins were transferred using the Trans-Blot Turbo system onto PVDF or nitrocellulose membranes Membranes were verified by Ponceau S staining and blocked in 5% milk in TBS-T.

The following primary antibodies were used at 1:1000: Phospho-Akt (Ser473) (D9E XP® Rabbit mAb #4060, Cell Signaling Technology), Phospho-S6 Ribosomal Protein (Ser240/244) (D68F8 XP® Rabbit mAb #5364, Cell Signaling Technology), Phospho-4E-BP1 (Thr37/46) (236B4, Rabbit mAb #2855, Cell Signaling Technology), Phospho-S6 Ribosomal Protein (Ser235/236) (#2211, Cell Signaling Technology), *α*-Tubulin Antibody (2144S, Cell Signaling Technology). Following incubation, membranes were washed and probed with anti-rabbit IgG, HRP-linked secondary antibody (#7074S, Cell Signaling Technology, RRID: AB 2099233) diluted 1:1000 in 5% milk in 1X TBS-T. Protein bands were visualized using SuperSignal West Pico (Pierce, #34580) using Imaging System Vilber Fusion Solos. Membranes were subsequently stripped using Pierce Restore PLUS Western Blot Stripping Buffer (Thermo Scientific) for re-probing. Western blot images were quantified using grayscale measurements in ImageJ software (NIH) normalized to the loading control.

### Quantification and statistical analysis

Flow cytometry data were analyzed using FlowJo 10 (BD Biosciences). Statistical Analyses were performed using Prism software (v.9 GraphPad) and results are represented as mean ±SEM, unless otherwise indicated. Comparisons for two groups were calculated using unpaired two-tailed Student’s t tests. Comparisons of more than two groups were calculated using two-way ANOVA with Tukey multiple comparison tests. We observed normal distribution and no difference in variance between groups in individual comparisons. Selection of sample size was based on extensive experience with metabolic assays. The number of independent experiments performed, and the p values for each experiment are reported in the corresponding figure legends.

## Notes

### Competing Interest Statement

The authors have declared no competing interest.

## REFERENCES

1. Clemente, J.C., Ursell, L.K., Parfrey, L.W., and Knight, R. (2012). The impact of the gut microbiota on human health: an integrative view. Cell 148, 1258–1270. 10.1016/j.cell.2012.01.035.

2. Hou, K., Wu, Z.X., Chen, X.Y., Wang, J.Q., Zhang, D., Xiao, C., Zhu, D., Koya, J.B., Wei, L., Li, J., and Chen, Z.S. (2022). Microbiota in health and diseases. Signal Transduct Target Ther 7, 135. 10.1038/s41392-022-00974-4.

3. Galkin, F., Mamoshina, P., Aliper, A., Lane, E., Moskalev, V., Gladyshev, V.N., and Zhavoronkov, A. (2020). Human Gut Microbiome Aging Clock Based on Taxonomic Profiling and Deep Learning. iScience 23, 101199. 10.1016/j.isci.2020.101199.

4. Litichevskiy, L., Considine, M., Gill, J., Shandar, V., Cox, T.O., Descamps, H.C., Wright, K.M., Amses, K.R., Dohnalova, L., Liou, M.J., et al. (2025). Gut metagenomes reveal interactions between dietary restriction, ageing and the microbiome in genetically diverse mice. Nat Microbiol 10, 1240–1257. 10.1038/s41564-025-01963-3.

5. Shen, X., Wang, C., Zhou, X., Zhou, W., Hornburg, D., Wu, S., and Snyder, M.P. (2024). Nonlinear dynamics of multi-omics profiles during human aging. Nat Aging 4, 1619–1634. 10.1038/s43587-024-00692-2.

6. Wilmanski, T., Gibbons, S.M., and Price, N.D. (2022). Healthy aging and the human gut microbiome: why we cannot just turn back the clock. Nat Aging 2, 869–871. 10.1038/s43587-022-00294-w.

7. Bana, B., and Cabreiro, F. (2019). The Microbiome and Aging. Annu Rev Genet 53, 239–261. 10.1146/annurev-genet-112618-043650.

8. Lopez-Otin, C., Blasco, M.A., Partridge, L., Serrano, M., and Kroemer, G. (2023). Hallmarks of aging: An expanding universe. Cell 186, 243–278. 10.1016/j.cell.2022.11.001.

9. The Prolongation of Life. (1908). Nature 77, 289–290. 10.1038/077289b0.

10. Clark, R.I., Salazar, A., Yamada, R., Fitz-Gibbon, S., Morselli, M., Alcaraz, J., Rana, A., Rera, M., Pellegrini, M., Ja, W.W., and Walker, D.W. (2015). Distinct Shifts in Microbiota Composition during Drosophila Aging Impair Intestinal Function and Drive Mortality. Cell Rep 12, 1656–1667. 10.1016/j.celrep.2015.08.004.

11. Sommer, F., Bernardes, J.P., Best, L., Sommer, N., Hamm, J., Messner, B., Lopez-Agudelo, V.A., Fazio, A., Marinos, G., Kadibalban, A.S., et al. (2025). Life-long microbiome rejuvenation improves intestinal barrier function and inflammaging in mice. Microbiome 13, 91. 10.1186/s40168-025-02089-8.

12. Thevaranjan, N., Puchta, A., Schulz, C., Naidoo, A., Szamosi, J.C., Verschoor, C.P., Loukov, D., Schenck, L.P., Jury, J., Foley, K.P., et al. (2017). Age-Associated Microbial Dysbiosis Promotes Intestinal Permeability, Systemic Inflammation, and Macrophage Dysfunction. Cell Host Microbe 21, 455–466 e454. 10.1016/j.chom.2017.03.002.

13. Barcena, C., Valdes-Mas, R., Mayoral, P., Garabaya, C., Durand, S., Rodriguez, F., Fernandez-Garcia, M.T., Salazar, N., Nogacka, A.M., Garatachea, N., et al. (2019). Healthspan and lifespan extension by fecal microbiota transplantation into progeroid mice. Nat Med 25, 1234–1242. 10.1038/s41591-019-0504-5.

14. Chen, S., Li, Q., Shi, H., Li, F., Duan, Y., and Guo, Q. (2024). New insights into the role of mitochondrial dynamics in oxidative stress-induced diseases. Biomed Pharmacother 178, 117084. 10.1016/j.biopha.2024.117084.

15. Han, B., Lin, C.J., Hu, G., and Wang, M.C. (2019). ’Inside Out’-a dialogue between mitochondria and bacteria. FEBS J 286, 630–641. 10.1111/febs.14692.

16. Martinez-Martinez, D., Peres, T.V., Gehling, K., Quintaneiro, L., Cabrera, C., Cherevatenko, M., Cutty, S.J., Best, L., Marinos, G., Zimmerman, J., et al. (2025). Chemotherapy modulation by a cancer-associated microbiota metabolite. Cell Syst 16, 101397. 10.1016/j.cels.2025.101397.

17. Pryor, R., Norvaisas, P., Marinos, G., Best, L., Thingholm, L.B., Quintaneiro, L.M., De Haes, W., Esser, D., Waschina, S., Lujan, C., et al. (2019). Host-Microbe-Drug-Nutrient Screen Identifies Bacterial Effectors of Metformin Therapy. Cell 178, 1299–1312 e1229. 10.1016/j.cell.2019.08.003.

18. Gupta, R., Durham, T.J., Chau, G., Kanai, M., Uddin, M.M., Lu, W., Argentieri, M.A., Karczewski, K.J., Howrigan, D., Natarajan, P., et al. (2026). Mechanism of age-related accumulation of mtDNA mutations in human blood. Nature. 10.1038/s41586-026-10569-6.

19. Trifunovic, A., Wredenberg, A., Falkenberg, M., Spelbrink, J.N., Rovio, A.T., Bruder, C.E., Bohlooly, Y.M., Gidlof, S., Oldfors, A., Wibom, R., et al. (2004). Premature ageing in mice expressing defective mitochondrial DNA polymerase. Nature 429, 417–423. 10.1038/nature02517.

20. Marti, J.M., Martinez-Martinez, D., Rubio, T., Gracia, C., Pena, M., Latorre, A., Moya, A., and C, P.G. (2017). Health and Disease Imprinted in the Time Variability of the Human Microbiome. mSystems 2. 10.1128/mSystems.00144-16.

21. Flor, S., Dost, T., Haase, M., Simon, R., Ederer, S., Kadibalban, A.S., Taubenheim, J., Olecka, M., Walker, A., Zimmermann, J., et al. (2025). Multi-omics analysis highlights the link of aging-related cognitive decline with systemic inflammation and alterations of tissue-maintenance. bioRxiv, 2025.2007.2013.662751. 10.1101/2025.07.13.662751.

22. Gabande-Rodriguez, E., Gomez de Las Heras, M.M., Ramirez-Ruiz de Erenchun, P., Simo, C., Garcia-Canas, V., Inohara, N., Berenguer-Lopez, I., Enriquez-Zarralanga, V., Fernandez-Almeida, A., Oller, J., et al. (2026). Butyrate extends health and lifespan in mice with mitochondrial deficiency. Nat Commun 17. 10.1038/s41467-026-70547-4.

23. Regina, A., Solagna, F., Estrada, M.S., Jacobs, M.M.E., Martinez-Martinez, D., Georgomanolis, T., Alibashikj, I., Gjurgji, S., Pearson, C., Chang, D., et al. (2026). Cardiolipin preserves T(reg) metabolic fitness and immune homeostasis in the gut. Nat Metab. 10.1038/s42255-026-01533-9.

24. Wang, Q., Xu, Z., Ding, X., Wang, A., Song, S., Zhang, S., Chen, Y., Ding, Y., Jiang, L., and Ding, X. (2025). Ten mouse organs proteome and metabolome atlas from adult to aging. Genome Med 17, 116. 10.1186/s13073-025-01535-4.

25. Joseph, G.A., Wang, S.X., Jacobs, C.E., Zhou, W., Kimble, G.C., Tse, H.W., Eash, J.K., Shavlakadze, T., and Glass, D.J. (2019). Partial Inhibition of mTORC1 in Aged Rats Counteracts the Decline in Muscle Mass and Reverses Molecular Signaling Associated with Sarcopenia. Mol Cell Biol 39. 10.1128/MCB.00141-19.

26. Xu, S., Zhao, G., Hanigan, M.D., Cantalapiedra-Hijar, G., and Li, M. (2025). Branched-chain amino acids in muscle growth: mechanisms, physiological functions, and applications. J Anim Sci Biotechnol 16, 164. 10.1186/s40104-025-01300-y.

27. Andreatta, M., Berenstein, A.J., and Carmona, S.J. (2022). scGate: marker-based purification of cell types from heterogeneous single-cell RNA-seq datasets. Bioinformatics 38, 2642–2644. 10.1093/bioinformatics/btac141.

28. Lei, Y., Guerra Martinez, C., Torres-Odio, S., Bell, S.L., Birdwell, C.E., Bryant, J.D., Tong, C.W., Watson, R.O., West, L.C., and West, A.P. (2021). Elevated type I interferon responses potentiate metabolic dysfunction, inflammation, and accelerated aging in mtDNA mutator mice. Sci Adv 7. 10.1126/sciadv.abe7548.

29. Smith, P., Willemsen, D., Popkes, M., Metge, F., Gandiwa, E., Reichard, M., and Valenzano, D.R. (2017). Regulation of life span by the gut microbiota in the short-lived African turquoise killifish. Elife 6. 10.7554/eLife.27014.

30. Kundu, P., Lee, H.U., Garcia-Perez, I., Tay, E.X.Y., Kim, H., Faylon, L.E., Martin, K.A., Purbojati, R., Drautz-Moses, D.I., Ghosh, S., et al. (2019). Neurogenesis and prolongevity signaling in young germ-free mice transplanted with the gut microbiota of old mice. Sci Transl Med 11. 10.1126/scitranslmed.aau4760.

31. Houghton, D., Stewart, C.J., Stamp, C., Nelson, A., Aj Ami, N.J., Petrosino, J.F., Wipat, A., Trenell, M.I., Turnbull, D.M., and Greaves, L.C. (2018). Impact of Age-Related Mitochondrial Dysfunction and Exercise on Intestinal Microbiota Composition. J Gerontol A Biol Sci Med Sci 73, 571–578. 10.1093/gerona/glx197.

32. Chuandong, Z., Hu, J., Li, J., Wu, Y., Wu, C., Lai, G., Shen, H., Wu, F., Tao, C., Liu, S., et al. (2024). Distribution and roles of Ligilactobacillus murinus in hosts. Microbiol Res 282, 127648. 10.1016/j.micres.2024.127648.

33. Gems, D. (2022). The hyperfunction theory: An emerging paradigm for the biology of aging. Ageing Res Rev 74, 101557. 10.1016/j.arr.2021.101557.

34. Schwarzer, M., Gautam, U.K., Makki, K., Lambert, A., Brabec, T., Joly, A., Srutkova, D., Poinsot, P., Novotna, T., Geoffroy, S., et al. (2023). Microbe-mediated intestinal NOD2 stimulation improves linear growth of undernourished infant mice. Science 379, 826–833. 10.1126/science.ade9767.

35. Shimizu, K., Muranaka, Y., Fujimura, R., Ishida, H., Tazume, S., and Shimamura, T. (1998). Normalization of reproductive function in germfree mice following bacterial contamination. Exp Anim 47, 151–158. 10.1538/expanim.47.151.

36. Munyoki, S.K., Goff, J.P., Reshke, A., Wilderoter, E., Mafarachisi, N., Kolobaric, A., Sheng, Y., Mullett, S.J., King, G.E., DeSchepper, J.D., et al. (2025). The microbiota extends the reproductive lifespan of mice by safeguarding the ovarian reserve. Cell Host Microbe 33, 1731–1747 e1738. 10.1016/j.chom.2025.09.006.

37. Lee, H.Y., Lee, S.H., Lee, J.H., Lee, W.J., and Min, K.J. (2019). The role of commensal microbes in the lifespan of Drosophila melanogaster. Aging (Albany NY) 11, 4611–4640. 10.18632/aging.102073.

38. Snyder, D.L., Pollard, M., Wostmann, B.S., and Luckert, P. (1990). Life span, morphology, and pathology of diet-restricted germ-free and conventional Lobund-Wistar rats. J Gerontol 45, B52–58. 10.1093/geronj/45.2.b52.

39. Tazume, S., Umehara, K., Matsuzawa, H., Yoshida, T., Hashimoto, K., and Sasaki, S. (1991). Immunological function of food-restricted germfree and specific pathogen-free mice. Jikken Dobutsu 40, 523–528. 10.1538/expanim1978.40.4_523.

40. Twilhaar, W.N., Chang, J.-C., Terooatea, T.W., Yabukami, H., Kato, S., Satoh-Takayama, N., Ahmadi, A., Hetherington, N., Jeyamohan, P., Kageyama, T., et al. (2025). Epigenomic and transcriptomic germ-free ageing atlas reveals sterile inflammation as an intrinsic ageing feature. bioRxiv, 2025.2011.2019.689100. 10.1101/2025.11.19.689100.

41. Storelli, G., Defaye, A., Erkosar, B., Hols, P., Royet, J., and Leulier, F. (2011). Lactobacillus plantarum promotes Drosophila systemic growth by modulating hormonal signals through TOR-dependent nutrient sensing. Cell Metab 14, 403–414. 10.1016/j.cmet.2011.07.012.

42. Cox, T.O., Devason, A.S., de Araujo, A., Mason, S., Subramanian, M., Salvador, A.F.M., Descamps, H.C., Kim, J., Zhu, Y., Litichevskiy, L., et al. (2026). Intestinal interoceptive dysfunction drives age-associated cognitive decline. Nature. 10.1038/s41586-026-10191-6.

43. rodents, F.w.g.o.r.o.g.f.h.m.o., rabbits, Mahler Convenor, M., Berard, M., Feinstein, R., Gallagher, A., Illgen-Wilcke, B., Pritchett-Corning, K., and Raspa, M. (2014). FELASA recommendations for the health monitoring of mouse, rat, hamster, guinea pig and rabbit colonies in breeding and experimental units. Lab Anim 48, 178–192. 10.1177/0023677213516312.

44. Ewels, P., Magnusson, M., Lundin, S., and Kaller, M. (2016). MultiQC: summarize analysis results for multiple tools and samples in a single report. Bioinformatics 32, 3047–3048. 10.1093/bioinformatics/btw354.

45. Bolger, A.M., Lohse, M., and Usadel, B. (2014). Trimmomatic: a flexible trimmer for Illumina sequence data. Bioinformatics 30, 2114–2120. 10.1093/bioinformatics/btu170.

46. Langmead, B., and Salzberg, S.L. (2012). Fast gapped-read alignment with Bowtie 2. Nat Methods 9, 357–359. 10.1038/nmeth.1923.

47. Blanco-Miguez, A., Beghini, F., Cumbo, F., McIver, L.J., Thompson, K.N., Zolfo, M., Manghi, P., Dubois, L., Huang, K.D., Thomas, A.M., et al. (2023). Extending and improving metagenomic taxonomic profiling with uncharacterized species using MetaPhlAn 4. Nat Biotechnol 41, 1633–1644. 10.1038/s41587-023-01688-w.

48. Cerda, M., Wheeler-Martin, K., Bruzelius, E., Ponicki, W., Gruenewald, P., Mauro, C., Crystal, S., Davis, C.S., Keyes, K., Hasin, D., et al. (2021). Spatiotemporal Analysis of the Association Between Pain Management Clinic Laws and Opioid Prescribing and Overdose Deaths. Am J Epidemiol 190, 2592–2603. 10.1093/aje/kwab192.

49. Mallick, H., Rahnavard, A., McIver, L.J., Ma, S., Zhang, Y., Nguyen, L.H., Tickle, T.L., Weingart, G., Ren, B., Schwager, E.H., et al. (2021). Multivariable association discovery in population-scale meta-omics studies. PLoS Comput Biol 17, e1009442. 10.1371/journal.pcbi.1009442.

50. Schwaiger, M., Rampler, E., Hermann, G., Miklos, W., Berger, W., and Koellensperger, G. (2017). Anion-Exchange Chromatography Coupled to High-Resolution Mass Spectrometry: A Powerful Tool for Merging Targeted and Non-targeted Metabolomics. Anal Chem 89, 7667–7674. 10.1021/acs.analchem.7b01624.

51. Demichev, V., Messner, C.B., Vernardis, S.I., Lilley, K.S., and Ralser, M. (2020). DIA-NN: neural networks and interference correction enable deep proteome coverage in high throughput. Nat Methods 17, 41–44. 10.1038/s41592-019-0638-x.

52. Tyanova, S., Temu, T., Sinitcyn, P., Carlson, A., Hein, M.Y., Geiger, T., Mann, M., and Cox, J. (2016). The Perseus computational platform for comprehensive analysis of (prote)omics data. Nat Methods 13, 731–740. 10.1038/nmeth.3901.

53. Hughes, C.S., Moggridge, S., Muller, T., Sorensen, P.H., Morin, G.B., and Krijgsveld, J. (2019). Single-pot, solid-phase-enhanced sample preparation for proteomics experiments. Nat Protoc 14, 68–85. 10.1038/s41596-018-0082-x.

